# DOT1L bridges transcription and heterochromatin formation at pericentromeres

**DOI:** 10.1101/2021.10.21.465349

**Authors:** Aushaq B. Malla, Haoming Yu, Srilekha Kadimi, TuKiet Lam, Andy L. Cox, Zachary D. Smith, Bluma J. Lesch

## Abstract

Repetitive DNA elements are packaged in heterochromatin, but many require bursts of transcription to initiate and maintain long-term silencing. The mechanisms by which these heterochromatic genome features are transcribed remain largely unknown. Here, we show that DOT1L, a conserved histone methyltransferase that modifies lysine 79 of histone H3 (H3K79), has a specialized role in transcription of major satellite repeats to maintain pericentromeric heterochromatin and genome stability. We discover that H3K79me3 is enriched at repetitive elements, that DOT1L loss specifically compromises pericentromeric satellite transcription, and that this function depends on interaction between DOT1L and the chromatin remodeler SMARCA5. Activation of pericentromeric repeats by DOT1L drives the first establishment of heterochromatin structures in cleavage-stage embryos and is required for preimplantation viability. Our findings uncover a vital instructive role for DOT1L as a bridge between transcriptional activation of heterochromatic repeats and maintenance of genome integrity, and illuminate global chromatin dynamics during early development.

## Introduction

Control of transcription by chromatin is central to metazoan development and frequently disrupted in disease. Most regions of the genome are packaged by either open, transcriptionally active euchromatin, or compact, transcriptionally repressed heterochromatin. Repeat sequences such as satellite elements and retrotransposons are classic examples of constitutively repressed heterochromatic elements. Paradoxically, RNA transcripts from repeat sequences are often required to establish or maintain the heterochromatic state, meaning that these regions must periodically accommodate active transcription (Camacho et al., 2018; Frescas et al., 2008; Fukagawa et al., 2004; Grewal and Jia, 2007; Zofall and Grewal, 2006). To date, most research on chromatin regulation at repetitive elements has focused on factors that set up and maintain transcriptional repression. A major unsolved problem is how coordinated pulses of transcriptional activity are enacted in the context of heterochromatin at repetitive sequences, and how this transcriptionally active state can be rapidly reversed to restore epigenetic silencing.

Chromatin composition and function is determined in part by covalent modifications to histone proteins, which modulate interactions between nucleosomes and chromatin binding factors (Hyun et al., 2017). DOT1L (Disruptor of telomeric silencing 1 like, also called KMT4) is an evolutionarily conserved histone methyltransferase that catalyzes mono-, di-, and tri-methylation of histone H3 at lysine (K) 79 (H3K79me1/2/3). DOT1L and H3K79 methylation are typically associated with transcriptional activation at single-copy genes (Guenther et al., 2008; Steger et al., 2008; Wood et al., 2018). H3K79me2 accumulates downstream of the promoter region at actively transcribed genes, and DOT1L interacts directly with the C-terminal domain of RNA Polymerase II (PolII) and with several PolII-associated elongation complexes (Jonkers et al., 2014; Kim et al., 2012; Mohan et al., 2010; Veloso et al., 2014; Wood et al., 2018). In leukemia, DOT1L interacts with elongation factors fused to the transcriptional coactivator MLL1 (mixed lineage leukemia 1) and promotes constitutive transcriptional activation of genes responsible for leukemogenesis (Bernt et al., 2011; Guenther et al., 2008).

However, other data suggest a contrasting role for DOT1L in heterochromatin regulation and formation. Loss of *Dot1* or mutation of the H3K79 residue impair telomeric silencing in budding yeast (Ng et al., 2002). In mouse embryonic stem cells (mESCs), *Dot1L* mutation reduces the constitutive heterochromatin modifications H3K9me2 and H4K20me3 at large arrays of tandem repeats that make up the bulk of centromeric and pericentromeric regions, with reciprocal gains in the euchromatin-associated modification H3K9ac (Jones et al., 2008).

Pericentromeric heterochromatin (PCH) is required for transposon suppression and chromosome segregation, and coordinates heterochromatin formation and maintenance across the genome (Janssen et al., 2018). PCH is composed of tandem arrays of specific repeating sequences called satellites; in mouse, this 234 base pair repeating sequence element is called the major satellite. Although packaged in heterochromatin and is generally silent, major satellite DNA is transcribed at specific times in the cell cycle and during development. Major satellite transcription is cell cycle dependent in mouse fibroblasts, with transcript levels peaking near the G1/S boundary and persisting through mitosis (Lu and Gilbert, 2007). In mESCs, transcription factors such as NANOG, SALL1, and YY1 are recruited to promoters within pericentromeric regions and may contribute to their transcriptional activity (Novo et al., 2016; Shestakova et al., 2004). In later development, major satellite transcripts accumulate in the mouse central nervous system from embryonic day 11.5 (E11.5) to E15.5, and are abundant in adult liver and testis (Rudert et al., 1995). Following fertilization, a sudden burst of major satellite transcription occurs beginning at the four-cell stage and is thought to be essential for reorganizing heterochromatin into densely stained structures called chromocenters prior to further developmental progression (Casanova et al., 2013; Probst et al., 2010). Transcriptional activity at major satellite sequences is proposed to recruit heterochromatin protein 1 (HP1), suggesting that major satellite transcription is essential for heterochromatinization at PCH (Frescas et al., 2008; Probst et al., 2010). In keeping with a requirement for appropriate major satellite regulation in maintaining normal cellular function, aberrant upregulation of satellite transcripts also occurs in human pathological states, including several cancers (Bersani et al., 2015; Ting et al., 2011).

Despite its importance in genome regulation and integrity, our understanding of how intermittent transcriptional activation is accomplished in heterochromatin elements such as PCH remains limited. Here, we reveal a critical role for DOT1L in stabilization and initiation of pericentromeric transcription in mESCs and mouse preimplantation embryos, respectively. We show that DOT1L selectively promotes transcriptional activity at major satellite elements, and establish the chromatin regulator SMARCA5 as a DOT1L partner that contributes to major satellite expression. DOT1L is required for major satellite transcription during preimplantation embryonic development, and its inhibition leads to cell cycle arrest and lethality in cleavage-stage embryos. Our work identifies DOT1L as a transcriptional activator at PCH and suggests that this activity is required for establishment of heterochromatin structure and embryo viability.

## Results

### H3K79me3 is enriched at repetitive elements in mouse embryonic stem cells

To understand the distribution of H3K79 methylation in mESC nuclei, we performed co-staining of H3K79me2 or H3K79me3 with markers for mitosis (H3S10P) and pericentromeric heterochromatin (HMGA1, High Mobility Group protein A1) (Jagannathan et al., 2018). The distribution of H3K79me2 is cell cycle dependent: it localized to chromosome arms during mitosis (**Figure 1A**), and to euchromatin in interphase cells (**Figure 1B**). In contrast, H3K79me3 strongly localized to HMGA1-enriched pericentromeric regions both in mitotic cells and at interphase (**Figure 1A, 1B**). These results suggest that H3K79me2 and H3K79me3 are differentially enriched within chromatin, with H3K79me2 accumulating in euchromatic and H3K79me3 in more heterochromatic regions, a conclusion supported by a similar observation in fibroblasts (Ooga et al., 2008).

**Figure 1.**
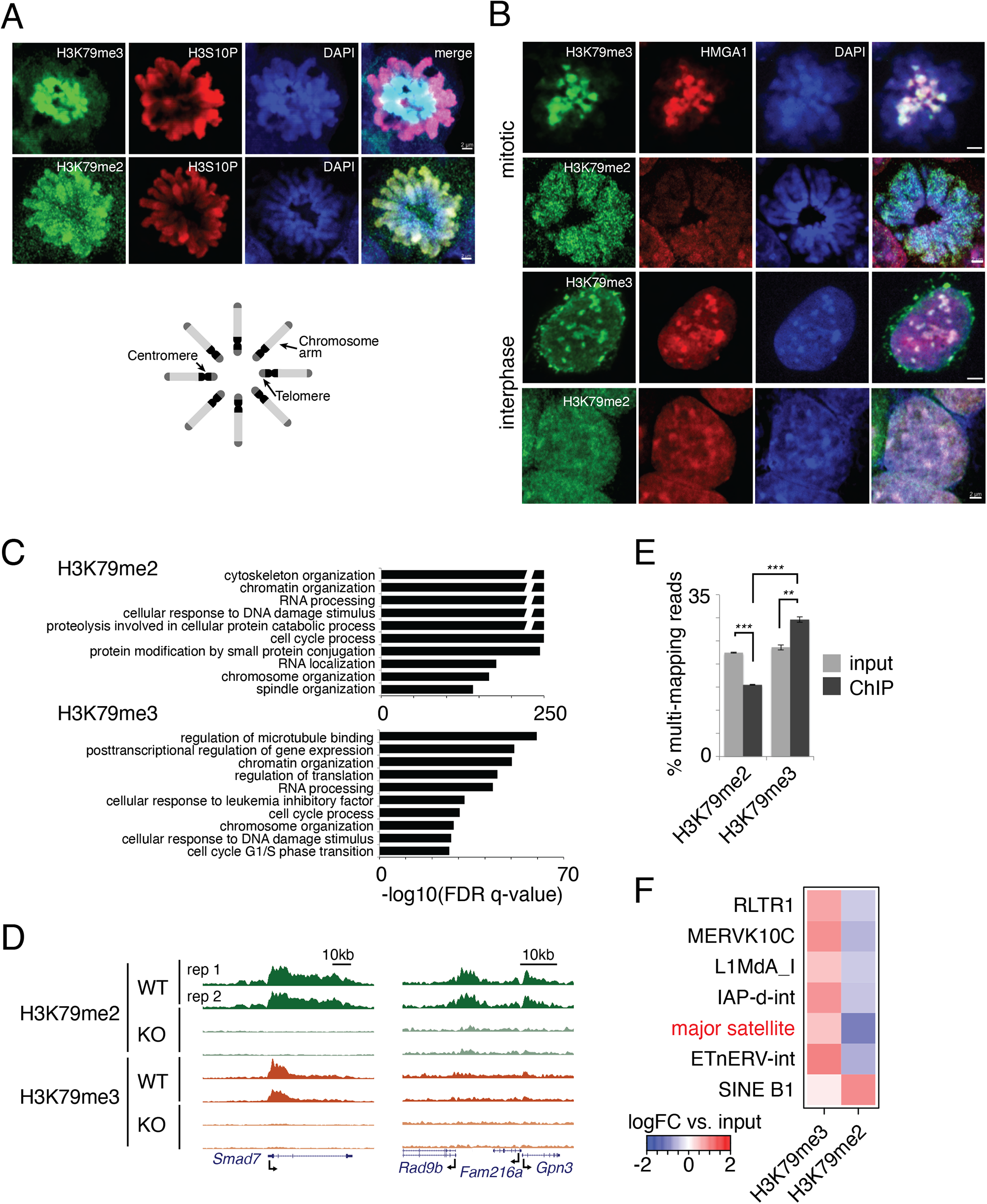
H3K79me3 distinguishes the repetitive genome from single-copy genes. (A) Immunofluorescence staining of single prometaphase nuclei showing H3K79me3 or H3K79me2 (green) and mitotic marker phospho-H3S10 (H3S10P, red). DNA is stained with DAPI (blue). Bottom, schematic of chromosomes at prometaphase showing arrangement of centromeres at the center of the metaphase plate. Scale bar, 2µm. (B) Immunofluorescence staining of single nuclei for H3K79me3 or H3K79me2 (green) and the pericentromeric heterochromatin marker HMGA1 (red). Upper panels, prometaphase; lower panels, interphase. DNA is stained with DAPI (blue). Scale bar, 2µm. (C) Selected gene ontology (GO) terms enriched among genes located near H3K79me2 or K3K79me3 peaks in mESCs. Break in the top 5 bars indicates a discontinuous axis. See Supplementary Table 2 for complete term lists. (D) Representative ChIP-seq tracks for H3K79me2 and H3K79me3 at a locus enriched for both marks (left) and a locus enriched for H3K79me2 only (right) in wild type and *Dot1L* KO mESCs. Two biological replicates are shown. (E) Fraction of ChIP and input libraries composed of non-uniquely mapping reads for H3K79me2 and H3K79me3 ChIP-seq data. **p<0.01, ***p<0.001, two-tailed Student’s t-test. (F) Heatmap of H3K79me2 and H3K79me3 ChIP-seq enrichment relative to input at selected repetitive element sequences.

To understand its genomic distribution in greater detail, we mapped both H3K79me2 and H3K79me3 in mESCs using chromatin immunoprecipitation followed by sequencing (ChIP-seq). To verify specificity of the ChIP antibody, we generated *Dot1L* knockout (KO) mESCs using CRISPR/Cas9 and confirmed loss of H3K79me2 and H3K79me3 by immunoblot and ChIP-seq in *Dot1L* KO cells (**Table S1**, **Figure S1**). We detected very few peaks (2 H3K79me2 and 67 H3K79me3 peaks) in *Dot1L* KO cells, validating our ChIP results. We eliminated these peaks as artifacts, defining a final set of 46,409 H3K79me2 peaks and 5,167 H3K79me3 peaks in mESCs. Enriched gene ontology (GO) categories for genes near H3K79me2 and H3K79me3 peaks were similar and consistent with known roles for DOT1L, including functions related to chromatin organization, cellular response to DNA damage, and cell cycle regulation (**Figure 1C**, **Table S2**).

Interestingly, careful analysis of ChIP-seq reads suggested that H3K79me3 was more likely than H3K79me2 to be enriched at repeat elements. At single-copy genes, many more loci were enriched for H3K79me2 than H3K79me3 (11,364 vs. 4,326 genes; **Table S2**, **Figure 1D**), and H3K79me2 signal was higher even at the subset of genes positive for both marks (**Figure S2A**, **S2B**). On the other hand, twice as many reads in our H3K79me3 libraries were non-uniquely mapped (mean 29.6% of reads for H3K79me3 compared to 15.5% for H3K79me2, p=0.0008, two-tailed Student’s t-test), indicating that they aligned to repetitive regions. This disparity was not apparent in non-immunoprecipitated input control libraries (mean 23.5% for H3K79me3 compared to 22.5% for H3K79me2). This finding implies that H3K79me3 is enriched in the repetitive genome. More formal analysis of enrichment at specific repetitive element consensus sequences using a dedicated analytic pipeline (Criscione et al., 2014), revealed that H3K79me3 was enriched (FDR q-value < 0.05) at 203 repeat elements in mESCs, whereas H3K79me2 was depleted (**Figures 1F**, **S2C**). We confirmed the relative enrichment of H3K79me3 compared to H3K79me2 at a subset of these elements by ChIP-qPCR (**Figure S2D**). These results suggest unique functions for H3K79me3 within the repetitive genome, including at major satellite repeats and retrotransposons.

### DOT1L is a transcriptional activator at major satellite repeats

The strong immunofluorescence-based localization of H3K79me3 to pericentromeric regions (**Figure 1A, 1B**) and enrichment for major satellite sequences in our ChIP-seq data (GSAT_MM, q = 4.52×10^-6^) prompted us to examine a potential role for DOT1L in regulation of pericentromeric heterochromatin. To see if major satellite expression was altered in the absence of DOT1L, we performed transcriptional profiling using ribosome-depletion RNA-seq in *Dot1L* KO mESCs (**Table S1**). Overall, DOT1L ablation has only a modest effect on protein-coding gene expression, with a total of 1,149 differentially expressed genes in *Dot1L* KO mESCs (log2 fold change ≥ 1, adjusted p-value ≤ 0.05) (**Table S3**). Our differentially expressed genes were not biased toward up- or downregulation (662 genes (58%) upregulated and 487 genes (42%) downregulated) (**Figure 2A**, **Table S3**). Almost half (45%) of downregulated genes are enriched for H3K79me2, compared to 19% for H3K79me3 (85% of which were also marked by H3K79me2) (**Figure S3A**), consistent with the known association between H3K79me2 and transcriptional elongation at single-copy genes (Guenther et al., 2008; Steger et al., 2008; Veloso et al., 2014). We did not detect any significant functional enrichment or obvious pattern in the set of downregulated genes, although upregulated genes were enriched for some pathways known to be associated with DOT1L function, such as developmental regulation and control of cell death (**Figure 2B**, **Table S3**). Together, these results support a role for DOT1L and H3K79 methylation, especially H3K79me2, in the regulation of single-copy genes, and confirm previous findings that loss of DOT1L has only modest effects on single-copy gene expresson in mESCs (Cao et al., 2020).

**Figure 2.**
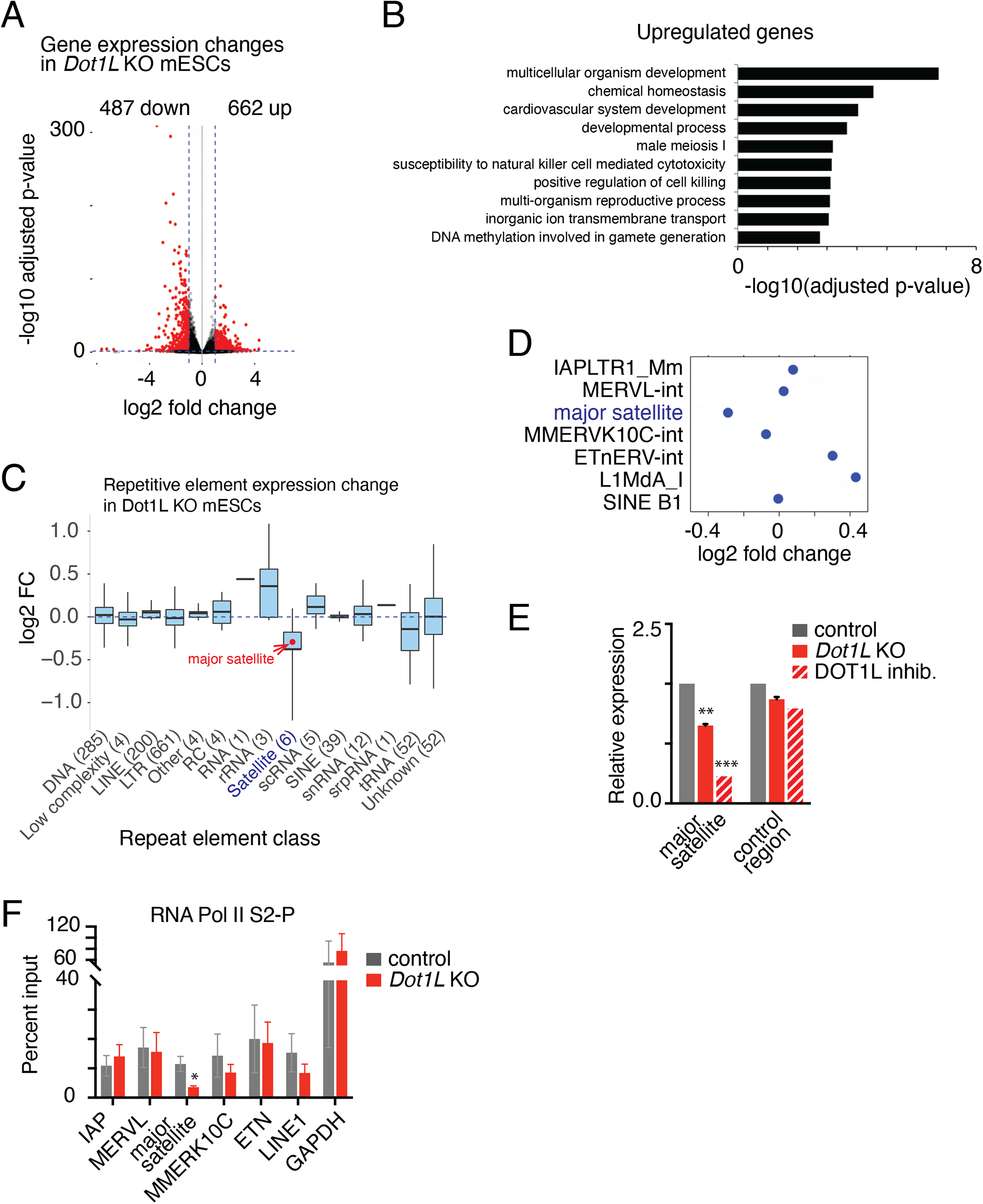
DOT1L promotes transcriptional activity at major satellites. (A) Volcano plot of differentially expressed single-copy genes in *Dot1L* KO mESCs (log2 fold change ≥ 1, adjusted p-value ≤ 0.05). Each point represents one gene. Differentially expressed genes are shown in red. (B) Selected GO terms enriched among significantly upregulated single-copy genes in *Dot1L* KO mESCs. See Supplementary Table 4 for complete term lists. (C) Differential expression of repeat element classes in *Dot1L* KO relative to control mESCs. Red dot indicates relative expression of major satellite transcripts within the broader set of annotated satellite sequences. Boxes represent interquartile range (25^th^-75^th^ percentile) of uniquely annotated repeat classes within each broad category; whiskers represent maximum and minimum values. Total number of repeat element annotations included in each class is shown in parentheses. (D) Strip plot showing relative expression of selected specific repeat element annotations in *Dot1L* KO mESCs compared to control. (E) Quantitative real-time PCR (RT-qPCR) of major satellite transcripts in wild type mESCs, *Dot1L* KO mESCs, and wild type mESCs treated with DOT1L inhibitor. Bars represent mean of four biological replicates for control and *Dot1L* KO, and two biological replicates for inhibitor-treated cells. Values were normalized to *Gapdh* as an internal control. Error bars represent ± standard error of the mean (SEM). **p < 0.001, ***p < 0.0001, two-tailed Student’s t test. (F) ChIP-qPCR for PolII-S2P at selected repeat elements. Bars represent mean of three biological replicates and error bars represent ± SEM. *p < 0.05, two-tailed Student’s t test.

When we examined expression of repetitive elements, we found that there was a general trend toward downregulation of satellite elements, including major satellites (**Figure 2C, 2D**). We confirmed this result with RT-qPCR in *Dot1L* KO mESCs and in wild type mESCs treated with the DOT1L inhibitor SGC0946 (Yu et al., 2012) (**Figure 2E**). In contrast, expression of other classes of repetitive elements, including LTRs, LINES, and SINEs, was either unchanged or slightly upregulated (**Figure 2C**, **2D**, **S3B**).

Downregulation of major satellite transcripts following *Dot1L* knockout or inhibition could be explained by either reduced transcriptional activity at these elements, or by altered posttranscriptional regulation. To distinguish between these possibilities, we performed ChIP-qPCR for RNA Polymerase II phosphorylated at Serine 2 of the C-terminal domain (PolII-S2P), the modified form of PolII associated with transcriptional elongation. We observed significant depletion of PolII-S2P at major satellites but not at other repetitive elements in *Dot1L* KO mESCs, consistent with a previously unknown requirement for DOT1L in driving transcription of major satellites (**Figure 2F**).

### DOT1L coordinates with SMARCA5 and heterochromatin factors to promote major satellite transcription at PCH

Major satellite transcripts derived from pericentromeric repeats play an essential role in the *de novo* reestablishment of heterochromatin after each mitotic division (Nair et al., 2020). In mice, major satellite transcripts themselves appear to recruit the H3K9 methyltransferase SUV39H1 to facilitate H3K9me3 deposition, and DOT1L has been shown to regulate deposition of heterochromatin marks at centromeres and telomeres (Jones et al., 2008; Shirai et al., 2017). Therefore, we predicted that DOT1L could promote heterochromatin formation at PCH through regulation of major satellite transcription. We performed ChIP-qPCR for the heterochromatin protein HP1β at major satellite repeats in wild type and *Dot1L* KO mESCs, and found that HP1β binding was specifically reduced at major satellites compared to other repetitive elements (**Figure 3A**), a result we confirmed by immunofluorescence (**Figure 3B**). These results suggest that DOT1L functions in maintaining pericentromeric heterochromatin through its regulation of major satellite transcription.

**Figure 3.**
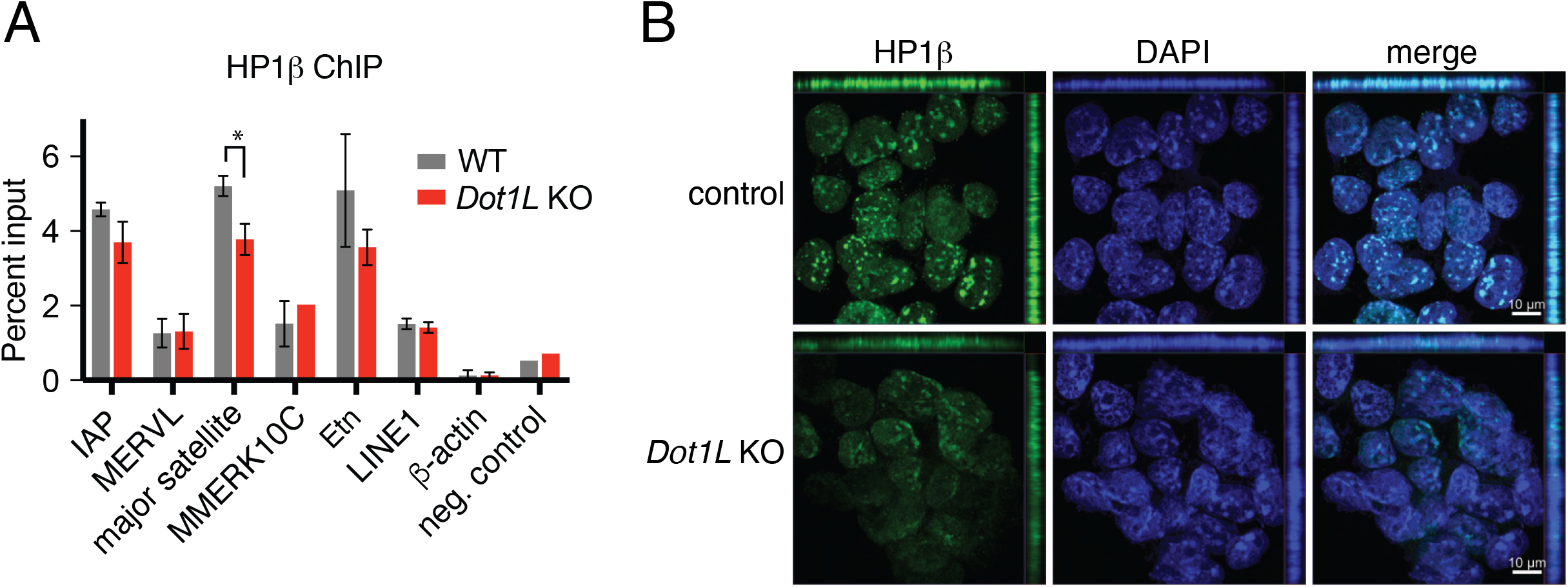
DOT1L stabilizes heterochromatin structure at PCH. (A) ChIP-qPCR for HP1β at selected repeat elements in *Dot1L* KO and wild type mESCs. Bars represent mean of two biological replicates and error bars represent ± SEM. *p < 0.05, two-tailed Student’s t test. (B) Immunofluorescence staining of HP1β (green) in wild type and *Dot1L* KO mESCs. DNA is stained with DAPI (blue). Z-axis signal is shown on the top and right of each image. Scale bar, 10µm.

The complex interaction between DOT1L, transcriptional activation, and heterochromatin reflects a longstanding challenge in understanding the molecular regulation of PCH. PCH fluctuates between transcriptionally active and heavily repressed states depending on the biological context, making it difficult to define straightforward regulatory pathways using genetic knockouts. To better define the function of DOT1L at PCH, we sought to discover the protein factors that recruit DOT1L to these regions. We overexpressed HA-tagged full-length DOT1L in mESCs and performed immunoprecipitation followed by mass spectrometry (IP-MS). A total of 70 protein interactors were identified (≥2 unique peptides) in transfected compared to non-transfected cells (**Table S4**). As expected, the top hit was MLLT10 (AF10), a transcriptional activator and known DOT1L interactor (**Figure 4A**) (Bitoun et al., 2007; Mohan et al., 2010). Overall, the list of interactors was enriched for pathways known to be associated with DOT1L function, including gene expression, DNA repair, chromatin remodeling, and histone modification (**Figure 4B**, **Table S4**).

**Figure 4.**
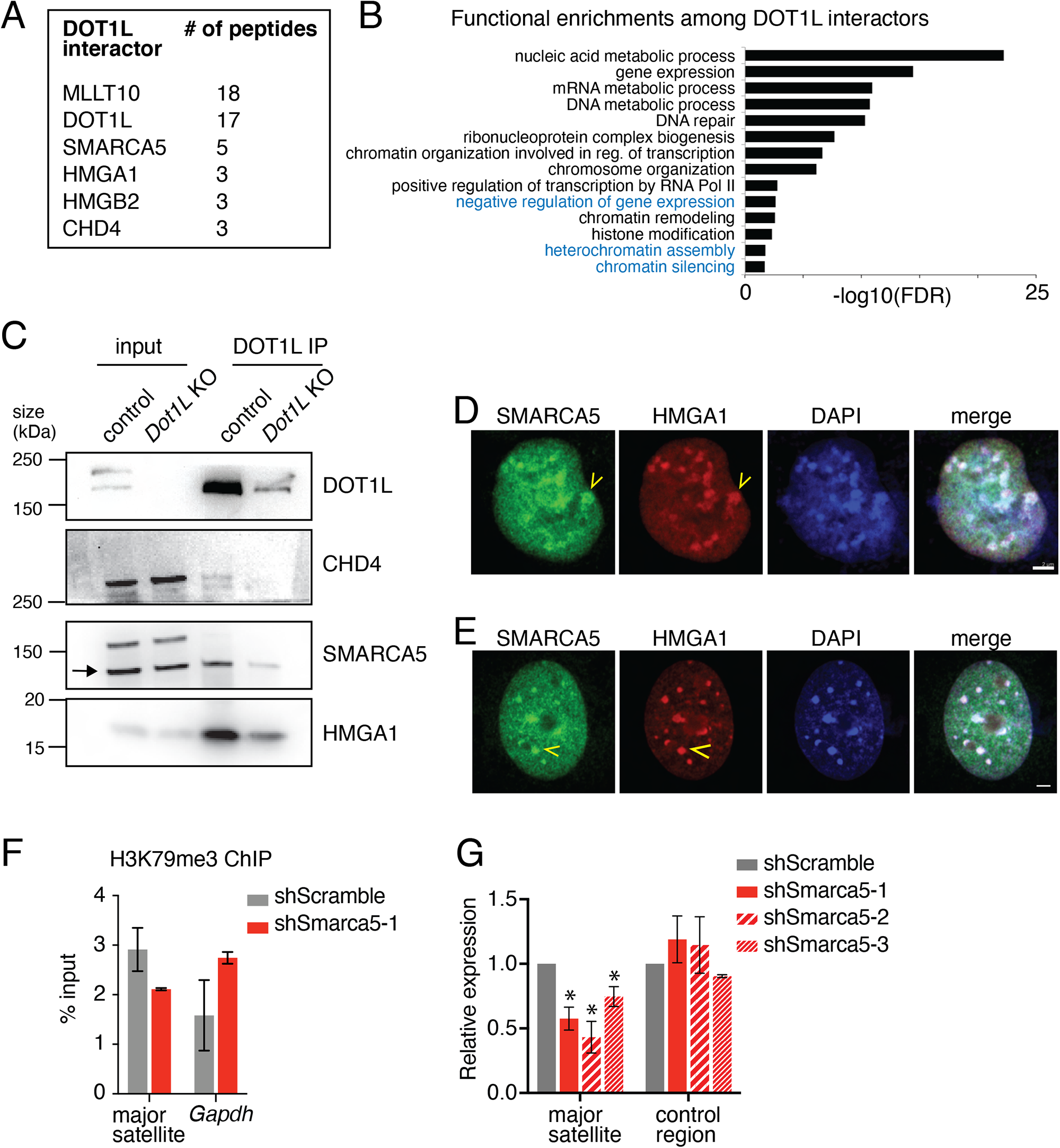
DOT1L interacts with SMARCA5 to regulate major satellite transcription at PCH. (A) Selected proteins detected by DOT1L IP-MS along with their unique peptide count. See Supplementary Table 5 for complete list of interactors. (B) Selected GO terms enriched among proteins that were co-immunoprecipitated with DOT1L. Terms associated with transcriptional repression are highlighted in blue. See Supplementary Table 6 for complete list of terms. (C) Western blot for candidate heterochromatin-associated interactors following co-immunoprecipitation with DOT1L. mESCs were grown on feeder cells, resulting in faint residual bands in the *Dot1L* KO lane. (D) Immunofluorescence staining of SMARCA5 (green) and HMGA1 (red) in mESCs, indicating localization of SMARCA5 to heterochromatic regions. Arrowheads indicate an example of SMARCA5 enrichment at PCH. DNA is stained with DAPI (blue). Scale bar, 4µm. (E) Immunofluorescence staining of SMARCA5 (green) and HMGA1 (red) in mouse fibroblasts, indicating localization of SMARCA5 to heterochromatic regions. Arrowheads indicate and example of SMARCA5 enrichment at PCH. DNA is stained with DAPI (blue). Scale bar, 4µm. (F) ChIP-qPCR for H3K79me3 at major satellite sequences in control and *Smarca5* knockdown (KD) mESCs. Bars represent mean of two biological replicates. Error bars represent represent ± SEM. (G) RT-qPCR for major satellite transcripts in control and *Smarca5* KD mESCs. Data represent transcript levels in three different *Smarca5* KD clones that stably express a single shRNA against *Smarca5*. Bars represent mean of two biological replicates. Error bars represent ± SEM. *p < 0.05, two-tailed Student’s t test.

Interestingly, there was also significant enrichment among the list of DOT1L interactors for pathways related to transcriptional repression, such as negative regulation of gene expression, heterochromatin assembly, and chromatin silencing (**Figure 4B**). Several of these interactors are known regulators of heterochromatin but have not previously been shown to interact with DOT1L. One such protein was HMGA1, a marker for PCH (Jagannathan et al., 2018), supporting the model that DOT1L is recruited to PCH to mediate major satellite transcription. Another intriguing interactor was SMARCA5 (SNF2H). SMARCA5 is the mammalian homolog of *Drosophila* ISWI, an ATP-dependent chromatin remodeler that promotes transcription by disrupting DNA-histone contacts to slide or evict histones (Aihara et al., 1998; Clapier and Cairns, 2009; Stopka et al., 2000), and has been reported to localize to PCH (Vargova et al., 2009). We validated the interactions between endogenous DOT1L and HMGA1, SMARCA5, and CHD4, another heterochromatin-associated factor, in wild type mESCs by co-immunoprecipitation, and confirmed that these interactions were abrogated in DOT1L KO mESCs (**Figure 4C**). We further confirmed localization of SMARCA5 to PCH by co-staining with HMGA1 in mESCs and fibroblasts (**Figure 4D**, **4E**).

We then asked if the factors that interact with DOT1L at PCH are required for the function of DOT1L in promoting major satellite expression. We chose SMARCA5 as a candidate target for further study. Complete knockout of SMARCA5 leads to apoptosis of inner cell mass cells (Stopka and Skoultchi, 2003), suggesting that *Smarca5* knockout mESCs are likely to be inviable. We therefore performed shRNA knockdown (KD) experiments to assess the role of SMARCA5 in major satellite expression. We recovered multiple *Smarca5* KD clones, with levels of SMARCA5 protein ranging from 1% to 40% of control (**Figure S4**). Consistent with our expectations, *Smarca5* KD cells exhibited slow growth and elevated apoptosis depending on the level of knockdown. H3K79me3 signal was reduced at major satellites in *Smarca5* KD cells compared to control (**Figure 4F**) and major satellite transcript levels were reduced by 50% (**Figure 4G**). Together, our results suggest that DOT1L coordinates with SMARCA5 to promote major satellite transcription and heterochromatinization at PCH.

### Loss of DOT1L leads to chromosomal abnormalities and cell death in mESCs

PCH stabilizes the centromeric core during mitosis (Yi et al., 2018), and transcription of major satellites has been proposed to reinforce heterochromatinization at PCH during anaphase, suggesting that it is required for mitotic progression (Lu and Gilbert, 2007; Saksouk et al., 2015). Since we find that DOT1L promotes major satellite expression, we predicted that DOT1L loss would result in defects in chromosome segregation during mitotic division. From our immunofluorescence data (**Figure 1A**), we observed that a fraction of Dot1L KO cells exhibited aberrant chromosome alignment at the metaphase plate resulting in disorganized chromosome congression (**Figure 5A**). To further characterize this defect, we examined metaphase chromosome spreads in *Dot1L* KO and control mESCs, and found that *Dot1L* KO cells had significantly higher levels of chromosome breakage (**Figure 5B** (arrowheads) and **5C**) as well as a trend toward higher frequency of premature centromere separation (**Figure 5D**). We did not observe a significant increase in chromosome fusions in *Dot1L* KO cells (**Figure 5E**). Collectively, these results demonstrate that DOT1L loss predisposes cells to chromosomal rearrangements and genomic instability, which may contribute to the cell proliferation defects previously reported in mESCs lacking DOT1L (Jones et al., 2008).

**Figure 5.**
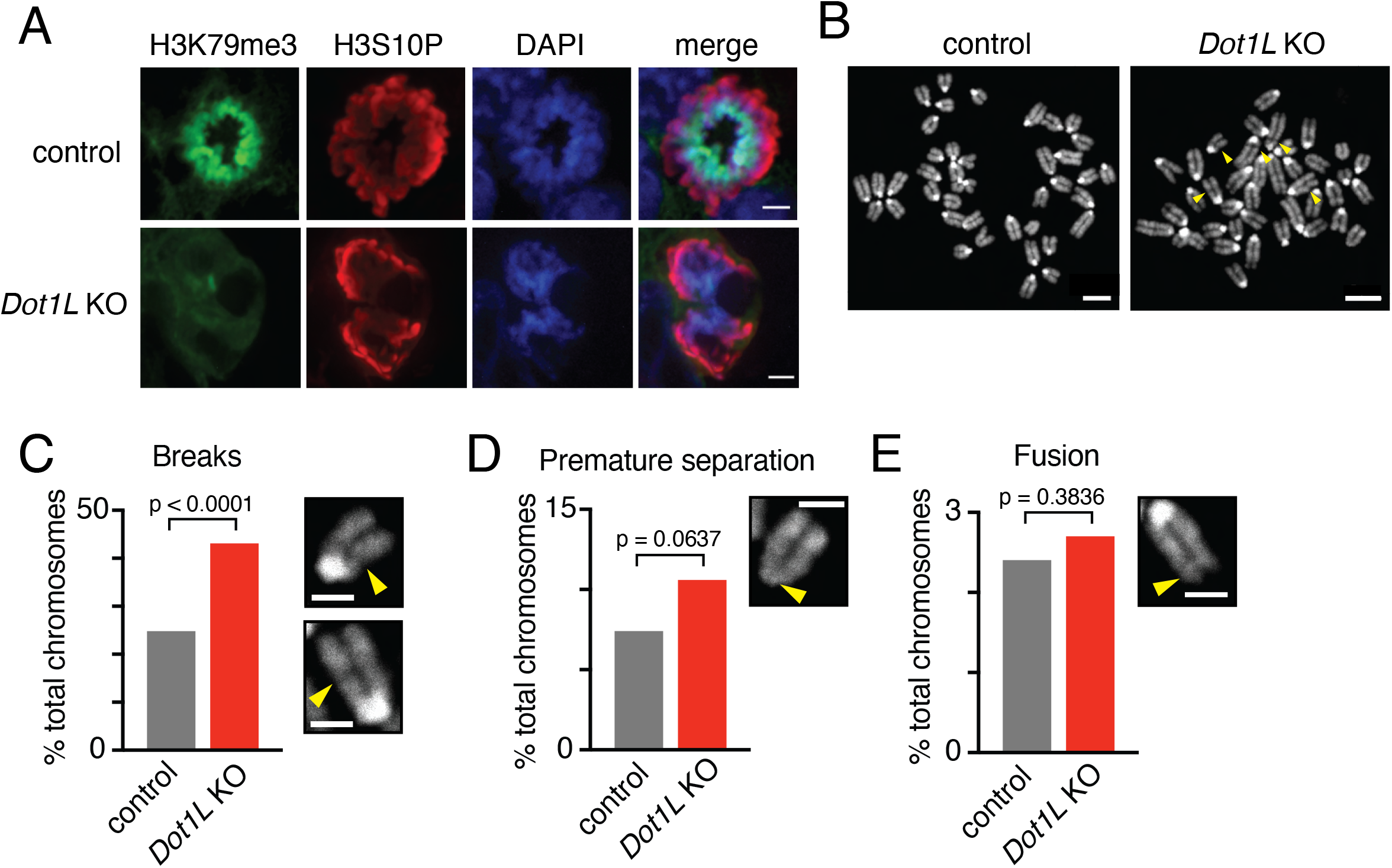
Loss of DOT1L causes mitotic defects and chromosomal abnormalities in mESCs. (A) Immunofluorescence staining of H3K79me3 (green) and H3S10P (red) in a single nucleus of control and *Dot1L* KO mESCs. DNA is stained with DAPI (blue). Scale bar, 5 µm. (B) Example metaphase spreads in wild type and *Dot1L* KO mESCs showing chromosome breaks (arrowheads). (C-E) Quantitation of chromosome defects in *Dot1L* KO mESCs, including chromosomal breaks and fragments (C), premature centromere separation (D), and chromosome fusions (E). Insets show representative examples of each chromosome defect. Scale bar, 2 µm. N=201 (wild type) or n=445 (KO) chromosomes from n=46 (wild type) or n=65 (KO) individual nuclei in two independent experiments. p-values, Fisher’s Exact test.

### DOT1L activity is required for cell cycle progression and viability in preimplantation embryos

We next asked whether the loss of DOT1L activity at PCH has *in vivo* significance. Major satellite expression at the four- to eight-cell stage in mouse preimplantation embryos is required for heterochromatin establishment and formation of chromocenters (Almouzni and Probst, 2011; Probst et al., 2010). We therefore asked whether DOT1L is required for upregulation of major satellite expression in early embryogenesis. First, we characterized the dynamics of H3K79me2 and H3K79me3 in preimplantation mouse embryos by immunostaining. For comparison, we co-stained with H3K9me3, which is present specifically in the maternal pronucleus in the zygote and localizes to heterochromatin as it forms in the four-cell stage (Burton and Torres-Padilla, 2010; Liu et al., 2004). H3K79me2 was absent from the zygote at least through at least the eight-cell stage (**Figure S5A**). In contrast, H3K79me3 was present at low levels in the early female pronucleus at 4 hours post fertilization (hpf), depleted in the late zygote by 10hpf, and reestablished at chromocenters as they form at the four-cell stage (48hpf) (**Figure 6A**). These results suggest that H3K79me2 and H3K79me3 are dynamically and differentially regulated during early stages of preimplantation development, and that DOT1L methyltransferase activity is present at nascent chromocenters by the four-cell stage.

**Figure 6.**
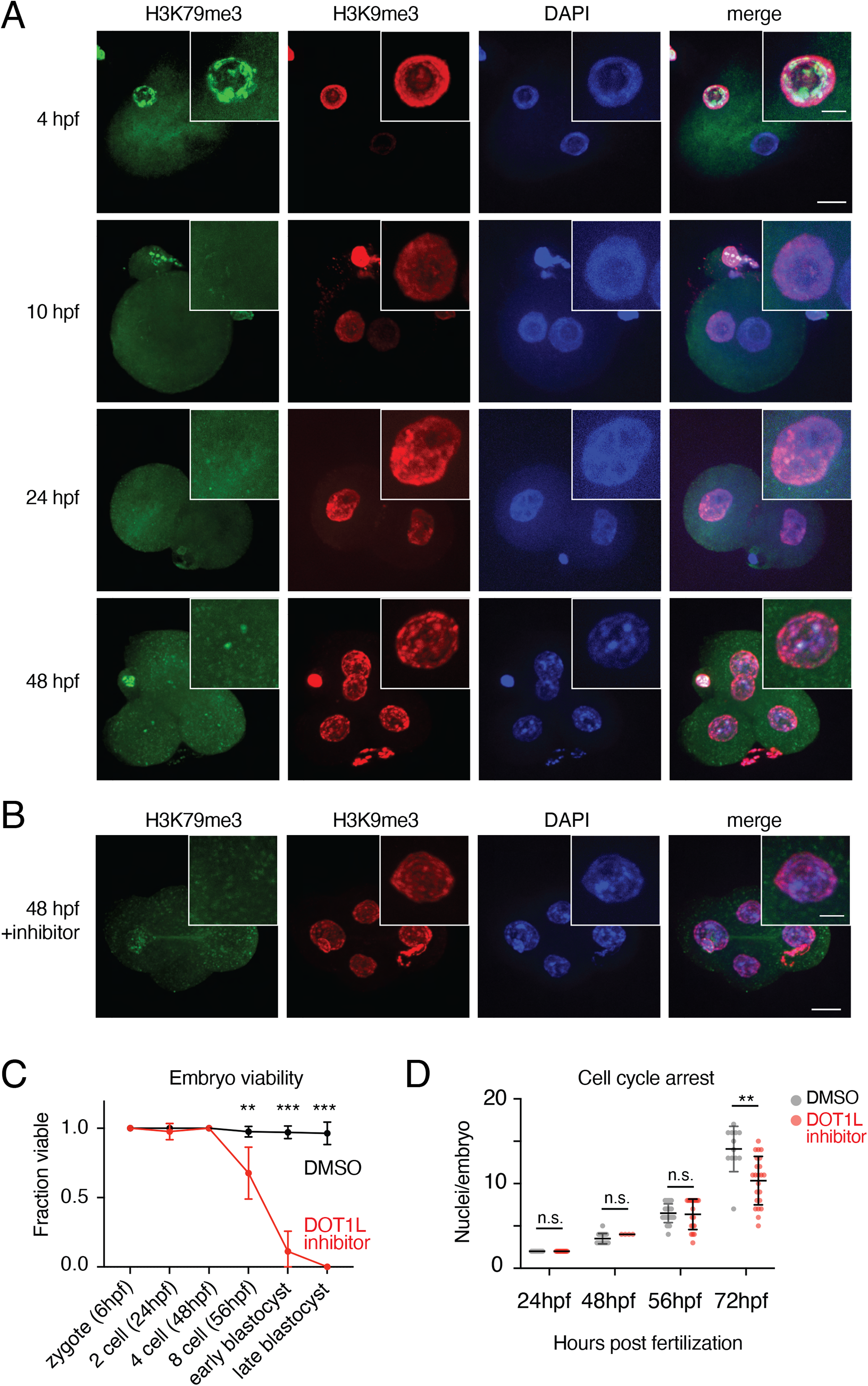
Dot1L activity is essential for cell cycle progression and viability during preimplantation development. (A) Immunofluorescence staining of H3K79me3 (green) and H3K9me3 (red) in wild type embryos during the first 48 hours of preimplantation development. DNA is stained with DAPI (blue). Scale bar, 10 µm (main image) or 4 µm (inset). (B) Immunofluorescence staining of H3K79me3 (green) and H3K9me3 (red) in DOT1L inhibitor-treated embryos at 48 hpf. DNA is stained with DAPI (blue). Scale bar, 10µm (main image) or 4 µm (inset). (C) Quantification of the fraction of fertilized zygotes that progress to a given state at each time point in control (vehicle treated) and inhibitor-treated embryos. Numbers of embryos that successfully progressed to a given stage were counted at each time point and the fraction for each is shown. Points represent the mean of 4-5 independent experiments. **q < 0.01, ***q < 0.001; false discovery rate (FDR) q-values calculated by unpaired t-test with two-state step-up correction for multiple comparisons. (D) Quantification of cell number per embryo at indicated time points in control (DMSO treated) and inhibitor-treated embryos. Numbers of nuclei per embryo were counted at each time point in three independent experiments. Bars represent mean and standard deviation. **q < 0.01; false discovery rate (FDR) q-values calculated by unpaired t-test with two-state step-up correction for multiple comparisons. n.s., not significant at a threshold of q < 0.05.

We next investigated whether DOT1L activity was required for preimplantation development by treating embryos *in vitro* with a small-molecule specific inhibitor of DOT1L (SGC0946). Fertilized zygotes were cultured in 10µM inhibitor and examined for up to 96 hours to evaluate the effect of DOT1L inhibition on embryonic progression to blastocyst. Immunostaining at intermediate time points confirmed that H3K79me3 was absent or substantially reduced in inhibitor-treated embryos (**Figure 6B**), although some embryos displayed residual staining that could reflect incomplete inhibition. In comparison to vehicle-treated embryos, DOT1L inhibition resulted in significant embryo death starting at 56hpf, with no embryos progressing past the early morula stage or to fully cavitated blastocysts (**Figure 6C**, **S5B**). While morphologically indistinguishable from controls in early cleavage, by 72hpf inhibitor-treated embryos failed to establish a cohesive and compact morphology typical of morula-stage embryos, suggesting pervasive developmental defects had accumulated prior to this initial differentiation event (**Figure S5C**).

To better understand the reason for the embryonic lethality, we counted the number of nuclei per embryo at each stage. Embryo progression through the third cleavage (eight cell stage) appeared identical between inhibitor and vehicle controls, but this was followed by a significant decrease in cell number by 72hpf, which was accompanied by an increase in the number of mitotic figures (**Figure 6D**). We also observed lagging chromosomes in anaphase cells of inhibitor-treated embryos beginning at the eight-cell stage (**Figure S5D**). These data suggest that mitotic dysfunction and progressive cell cycle arrest begin after the induction of major satellite transcription during late cleavage. Together with immunostaining data indicating that DOT1L enzymatic activity begins by 48hpf (**Figure 6A**), these results support a requirement for DOT1L in cell cycle progression beginning at the four- to eight-cell stage of preimplantation development.

### Inhibition of DOT1L activity in preimplantation embryos specifically downregulates major satellite transcription

A burst of major satellite transcription takes place in mouse embryos at the four-cell stage (Burton and Torres-Padilla, 2014), directly preceding the effects of DOT1L inhibition (**Figure 6C**). Given our finding that DOT1L promotes major satellite transcription at PCH in mESCs (**Figure 2**), we asked if DOT1L inhibition impairs major satellite transcription in preimplantation embryos. We performed ribosome depletion RNA-seq in eight-cell stage (56hpf) embryos cultured in the presence or absence of DOT1L inhibitor and assessed transcription of single-copy genes and repetitive elements (**Table S1**). We detected limited changes in expression of euchromatic single-copy genes following DOT1L inhibition: in total, 125 genes exhibit modest but statistically significant changes in expression in inhibitor-treated embryos (adjusted p-value < 0.05) (**Figure 7A**, **Table S5**), with a slight bias toward decreased expression as we saw in *Dot1L* KO mESCs (**Figure 7A**). Downregulated genes were significantly enriched for transcriptional and developmental functions, while upregulated genes were associated with elevated cell stress and activation of apoptotic pathways that could reflect emerging secondary effects (**Figure 7B**, **Table S5**).

**Figure 7.**
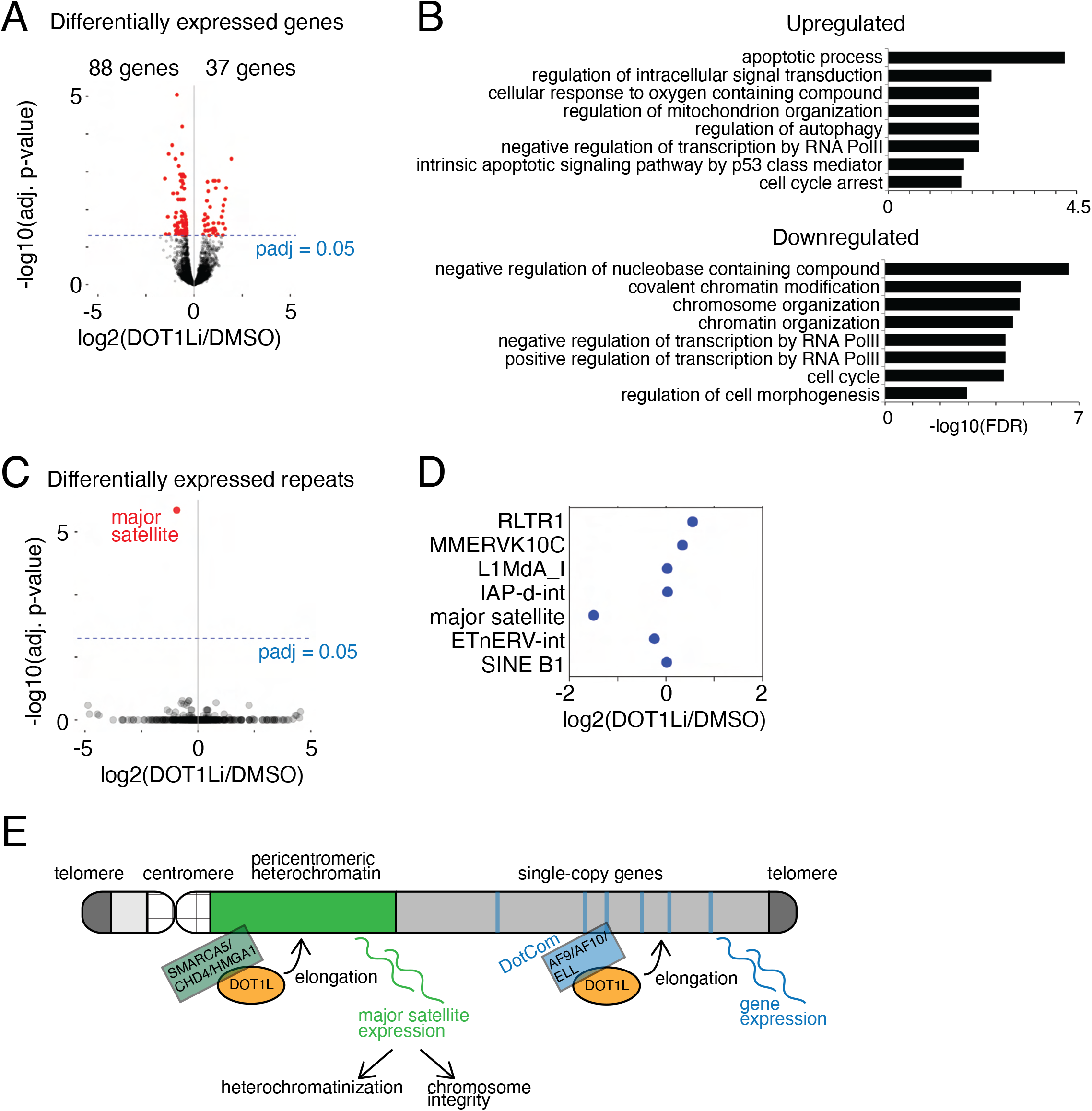
DOT1L promotes major satellite transcription in preimplantation embryos. (A) Volcano plot showing differentially expressed single-copy genes in DOT1L inhibitor-treated vs control embryos at 56hpf (8 cell stage). Each dot signifies one gene. Differentially expressed genes (adjusted p-value ≤ 0.05) are shown in red. (B) Selected enriched GO terms among significantly upregulated and downregulated single-copy genes in inhibitor-treated vs vehicle-treated embryos. (C) Volcano plot of annotated repeat elements in DOT1L inhibitor-treated vs vehicle-treated eight-cell embryos (56 hpf). Each dot signifies one annotated repetitive element category as indicated by RepeatMasker (Smit et al). Differentially expressed repetitive elements (adjusted p-value ≤ 0.05) are shown in red. The major satellite element is indicated. (D) Strip plot showing log2 fold change of selected repeat elements in eight-cell embryos as shown in C. (E) Model for DOT1L function at PCH and euchromatic single-copy genes. DOT1L coordinates with SMARCA5, CHD4, and HMGA1 for its recruitment to PCH to promote major satellite transcription, but interacts with the DotCom complex, including AF9, AF10, and ELL, at transcribed single copy genes.

Notably, among repetitive elements, we detected significant changes exclusively for major satellite transcripts (**Figure 7C, 7D**). Major satellite transcripts were significantly downregulated following drug treatment (log2 fold change -1.50, false discovery rate q-value 0.000450). This timing corresponds to localization of H3K79me3 to chromocenters in control embryos (**Figure 6A**) as well as to the onset of cell cycle defects observed in inhibitor-treated embryos (**Figure 6C**). Increased major satellite transcription at this time point may therefore reflect a critical moment in PCH establishment, which is subsequently required for cell cycle progression and embryo viability. Our results indicate that DOT1L activity is essential for this function.

## Discussion

In this study, we uncover a critical role for DOT1L in transcriptional activation of major satellites at pericentromeric heterochromatin, and find that this function is important for establishment of heterochromatin in preimplantation embryos and for heterochromatin maintenance and cell cycle progression in ESCs. While DOT1L is well known to be involved in transcriptional elongation at single-copy genes, transcriptional activation of major satellite repeats by DOT1L in PCH has not previously been reported and represents a novel regulatory role with significant biological impact.

We show that the DOT1L substrates H3K79me2 and H3K79me3 are differentially distributed within the repetitive genome. H3K79me3 is preferentially enriched at major satellites, and DOT1L is essential for promoting major satellite transcription. Notably, DOT1L is an evolutionarily ancient and unique histone methyltransferase with no cognate demethylase. How H3K79me3 may be read independently of H3K79me2 to signal downstream functions remains unclear, but could represent a novel signaling axis and bridge between transcription and heterochromatin formation at pericentromeric repeats. We propose that DOT1L participates in two distinct functional complexes: one that supports transcriptional elongation at single-copy genes and preferentially deposits H3K79me2 (Mohan et al., 2010), and a second, previously unknown complex that includes the repressive histone chaperones SMARCA5 and CHD4, operates to open and bind chromatin at PCH in a highly controlled fashion, and preferentially drives complete methylation to produce H3K79me3 (**Figure 7E**). Interestingly, DOT1L was recently reported to interact with other SWI/SNF chromatin remodelers, including SMARCA4, SMARCD2, SMARCC2, in K562 cells (Wu et al., 2021), and levels of H3K79 methylation were reduced in spermatogenic cells of mice heterozygous for a *Smarca5* knockout allele (Vargova et al., 2009), supporting a functional interaction between DOT1L and SMARCA5. Most previous work on the regulation of PCH and other repetitive heterochromatic elements has focused on the roles of canonical repressors. Our work highlights DOT1L as one of the first epigenetic regulators associated with transcriptional induction that appears essential to proper PCH initiation and maintenance. Notably, many of our results in DOT1L mutants appear to phenocopy the loss of repressive factors, such as the mitotic instability observed following loss of HP1 and in *Suv39h1/2* double knockout embryos (Abe et al., 2016; Peters et al., 2001).

Sequential conversion of H3K79me1 to H3K79me2 and H3K79me3 correlates with enhanced rates of transcription (Steger et al., 2008). Recently, an analysis of *Dot1L* KO mESCs found minimal impact on expression and no change in PolII-mediated elongation at single-copy genes, which are predominantly associated with H3K79me2 (Cao et al., 2020). Our data confirm minimal effects on protein-coding gene expression, but find a specific impact of *Dot1L* deletion on PolII elongation within major satellites, which are predominantly associated with H3K79me3. Together, these findings may imply a stronger effect of DOT1L on transcriptional elongation at PCH compared to euchromatic genes, an intriguing possibility to be explored in future work. We also found a slight preference for transcriptional upregulation (58% of differentially expressed genes) among single-copy genes following loss of DOT1L, which could indicate either a more widespread repressive role, as has been recently reported in hematopoietic progenitor cells (Borosha et al., 2020), or the emergence of secondary transcriptional effects.

We observed significant biological consequences of DOT1L disruption at PCH in both mESCs and early embryos. mESCs lacking DOT1L exhibited mitotic defects and increased chromosome damage at metaphase. During mitosis, chromosomes condense, and the kinetochore is assembled on centromeric core chromatin. During this process, PCH stabilizes the centromeric core (Yi et al., 2018). Transcriptional activity at major satellites has been proposed to reinforce heterochromatinization at PCH (Frescas et al., 2008) and stabilize mitotic progression (Lu and Gilbert, 2007; Saksouk et al., 2015). The increased frequency of mitotic defects in *Dot1L* KO mESCs, along with reduced recruitment of HP1β (an important structural component of heterochromatin that is implicated in cohesin recruitment to centromeres (Hahn et al., 2013)), suggests that DOT1L-mediated major satellite transcription is a key factor to maintain proper PCH structure and promote mitotic progression.

Excitingly, we identified DOT1L as a critical regulator of early development, where inhibition of DOT1L activity caused embryo arrest and death at preimplantation stages. In embryos, we found that H3K79me3 is enriched in chromocenters at the four-cell stage, when somatic heterochromatin structures, including chromocenters, first become prominent and major satellite expression is upregulated (Almouzni and Probst, 2011). Consistent with the temporal correlation between the appearance of H3K79me3 at chromocenters and the increase in major satellite transcripts, we found that DOT1L promotes major satellite transcription during early preimplantation development. Major satellite transcripts and DOT1L are both independently known to regulate mitotic progression (Biscotti et al., 2015; Kim et al., 2014; Müller and Almouzni, 2017), and the severe lethality phenotype in embryos following DOT1L inhibition could be the cumulative effect of reduced major satellite transcription on chromosome condensation, alignment, and segregation. Indeed, our RNA-seq data revealed upregulation of cell cycle and apoptotic pathways following DOT1L inhibition in 8-cell embryos. The delay between the initial upregulation of major satellite transcription at the two- to four-cell stage (Burton and Torres-Padilla, 2014; Casanova et al., 2013; Probst et al., 2010), and the phenotype we observe after the third cleavage, is also consistent with a mitotic defect that emerges from failed interphase regulation. Pericentromeric transcription occurs at the end of the G1 phase of the cell cycle (Lu and Gilbert, 2007), but the phenotype is delayed to the next mitosis. We propose that the cell cycle defects observed in DOT1L inhibitor-treated embryos are mediated by defective PCH establishment in the absence of major satellite transcription.

Notably, there is a discrepancy between the effects of DOT1L inhibition we report in preimplantation embryos and the phenotype of *Dot1L* zygotic null embryos, which survive until E10.5 (Jones et al., 2008). It is possible that maternal DOT1L protein from the oocyte is sufficient to establish PCH during preimplantation development and that this signature is maintained epigenetically in knockout embryos. In our experiments, maternal DOT1L would also be inhibited, effectively mirroring a combined maternal and zygotic disruption. Our RNA-seq data in DOT1L embryos was generated from F1 offspring of polymorphic mouse strains (see Methods), allowing us to distinguish paternal from maternal transcripts, and we found no bias in parent of origin for *Dot1L* transcripts; however, maternally inherited protein may still contribute to preimplantation epigenetic state. Recently, a maternal *Dot1L* conditional knockout was found to be dispensable for mouse development (Liao and Szabó, 2020), but a combined maternal-zygotic knockout has not been evaluated.

In addition to early embryogenesis, the role of DOT1L in major satellite expression may have *in vivo* significance in cancer. Several cancers are associated with DOT1L upregulation, including leukemia, lung adenocarcinoma, and colorectal cancer, and are also prone to overexpression of major satellite transcripts (Bersani et al., 2015; Ting et al., 2011). Heightened major satellite transcription due to DOT1L upregulation may contribute to tumor severity or progression in these cancers. Notably, SMARCA5 has also been implicated in pathogenesis of acute myeloid leukemia (AML), and like DOT1L is required for hematopoiesis (Kokavec et al., 2017; Stopka and Skoultchi, 2003; Zikmund et al., 2020). Future work will be important to better understand the potential role of DOT1L-mediated regulation of major satellite transcription in cancer.

In summary, our study defines a role for the H3K79 methyltransferase DOT1L in major satellite transcription and PCH regulation, illuminating a new mechanism for transient transcriptional activation in heterochromatin and uncovering the events that govern first establishment of heterochromatin in the early embryo. Future studies investigating this new function will be important for understanding the fundamental molecular mechanisms controlling heterochromatin dynamics across the cell cycle and throughout development.

## Supporting information

Table S1: sequencing summary

Table S2: ChIP

Table S3: mESC RNA-seq

Table S5: embryo RNA-seq

## Acknowledgments

We thank J. Wang and S. Guo for helpful discussions. We greatly appreciate help from the Yale Center for Genome Analysis for high-throughput sequencing. Proteomics data were collected on a mass spectrometer supported by NIH SIG S10OD018034 and Yale School of Medicine. We acknowledge start-up funds from Yale University School of Medicine and funding from the National Institute of Child Health and Human Development to B.J.L (R01HD098128), and support from the Searle Scholars Program to B.J.L. Bluma J. Lesch, M.D., Ph.D., holds a Career Award for Medical Scientists from the Burroughs Wellcome Fund.

## Author contributions

Conceptualization, A.B.M. and B.J.L; Validation, A.B.M. and H.Y.; Formal Analysis, A.B.M., H.Y., S.K., and B.J.L.; Investigation, A.B.M., H.Y., S.K., T.L., and Z.D.S.; Resources, Z.D.S. and B.J.L.; Writing – Original Draft, A.B.M.; Writing – Reviewing & Editing, A.L.C., Z.D.S, and B.J.L., Visualization, B.J.L.; Supervision, B.J.L.; Funding Acquisition, B.J.L.

## STAR Methods

### RESOURCE AVAILABILITY

#### Lead contact

Further information and requests for resources and reagents should be directed to and will be fulfilled by the lead contact, Bluma Lesch.

#### Materials availability

Plasmids and cell lines generated in this study are available from the lead contact upon request.

#### Data and code availability

RNA-seq and ChIP-seq datasets have been deposited at GEO and are publically available as of the date of publication under accession number GSE182744. Mass spectrometry datasets have been deposited at the ProteomeXchange Consortium via the PRIDE respository and are publically available as of the date of publication under accession number PXD028824. Original Western blot images, raw microscopy data, and raw qPCR data reported in this paper are available from the lead contact upon request.

This paper does not report original code.

Any additional information required to reanalyze the data reported in this paper is available from the lead contact upon request.

### EXPERIMENTAL MODEL AND SUBJECT DETAILS

#### Cell lines

##### Mouse embryonic stem cells (mESCs)

V6.5 male mouse embryonic stem cells (RRID:CVCL_C865) were grown in complete DMEM (Gibco, 11965-092) supplemented with 15% fetal bovine serum (Sigma-Aldrich, F2442), 1:100 pen-strep (Gibco, 15140-122), 1:100 GlutaMax (Gibco, 35050-061), 1:100 MEM NEAA (Gibco, 11140-050), Sodium Pyruvate 1:100 (Gibco, 11360-070), HEPES (Gibco, 15630-080), β mercaptoethanol (Sigma-Aldrich ,M6250), and 1:10000 LIF (Millipore, ESG1106). Cells were grown on a monolayer of mitotically arrested mouse embryonic fibroblasts in plates coated with 0.1% gelatin, maintained at 5% CO_2_ and passaged when they reached 70 to 80% confluence. Media was changed daily. No additional validation of the cell line was performed.

##### Mouse fibroblasts

NIH/3T3 cells (ATCC #CRL-1658, RRID:CVCL_0594) were grown in Dulbecco’s Modified Eagle’s Medium (DMEM, Life Technologies) supplemented with 10% fetal bovine serum (Life Technologies), 1 mM L-glutamine (Life Technologies), 100 U/ml penicillin, and 100 µg/ml streptomycin (Life Technologies). Cells were grown at 37°C at 5% CO_2_, and passaged when they reached 70-80% confluence. No additional validation of the cell line was performed.

##### Mice

All mice used in these studies were maintained and euthanized according to the principles and procedures described in the National Institutes of Health Guide for the Care and Use of Laboratory Animals. These studies were approved by the Yale University Institutional Animal Care and Use Committee under protocols 2020-20169 and 2020-20357 and conducted in accordance with the specific guidelines and standards of the Society for the Study of Reproduction. B6D2F1 strain (RRID:IMSR_JAX:100006) female mice (age 6–8 weeks, Jackson Labs) were superovulated for oocyte collection and zygotes were generated by ICSI using thawed CAST/EiJ strain sperm (RRID:IMSR_JAX:000928). For natural mating, superovulated B6D2F1 females were crossed with B6D2F1 males and zygotes isolated the following morning.

### METHOD DETAILS

#### Antibodies

| Antibody | Cat. No. | Application | Dilution |
| --- | --- | --- | --- |
| H3K79me1 | Abcam #ab2886 | WB | 1:1000 |
| H3K79me2 | Abcam #ab3594 | WB<br>IF<br>ChIP | 1:1000<br>1:500<br>1 ug/ 10<sup>6</sup> cells |
| H3K79me3 | Abcam #ab2621 | WB<br>IF<br>ChIP | 1:1000<br>1:500<br>1 ug/ 10<sup>6</sup> cells |
| DOT1L | Abcam #ab64077 | WB<br>IF | 1:400<br>1:250 |
| H3S10P | Abcam #ab14955 | IF | 1:500 |
| CHD4 | Abcam #ab70469 | WB | 1:500 |
| HMGA1 | Santa Cruz<br>#sc-393213 | IF | 1:200 |
| SMARCA5 | Thermo scientific<br># PA5-52601 | WB<br>IF | 1:1000<br>1:500 |
| RNA PolII S2P | Abcam # ab5095 | ChIP | 1 ug/ 10<sup>6</sup> cells |
| HP1 beta | CST # 2613 | IF<br>ChIP | 1:400<br>2 ug/ 10<sup>6</sup> cells |
| H3K9me3 | Novus Biologicals<br>#NBP1-30141 | IF | 1:500 |

#### Primers

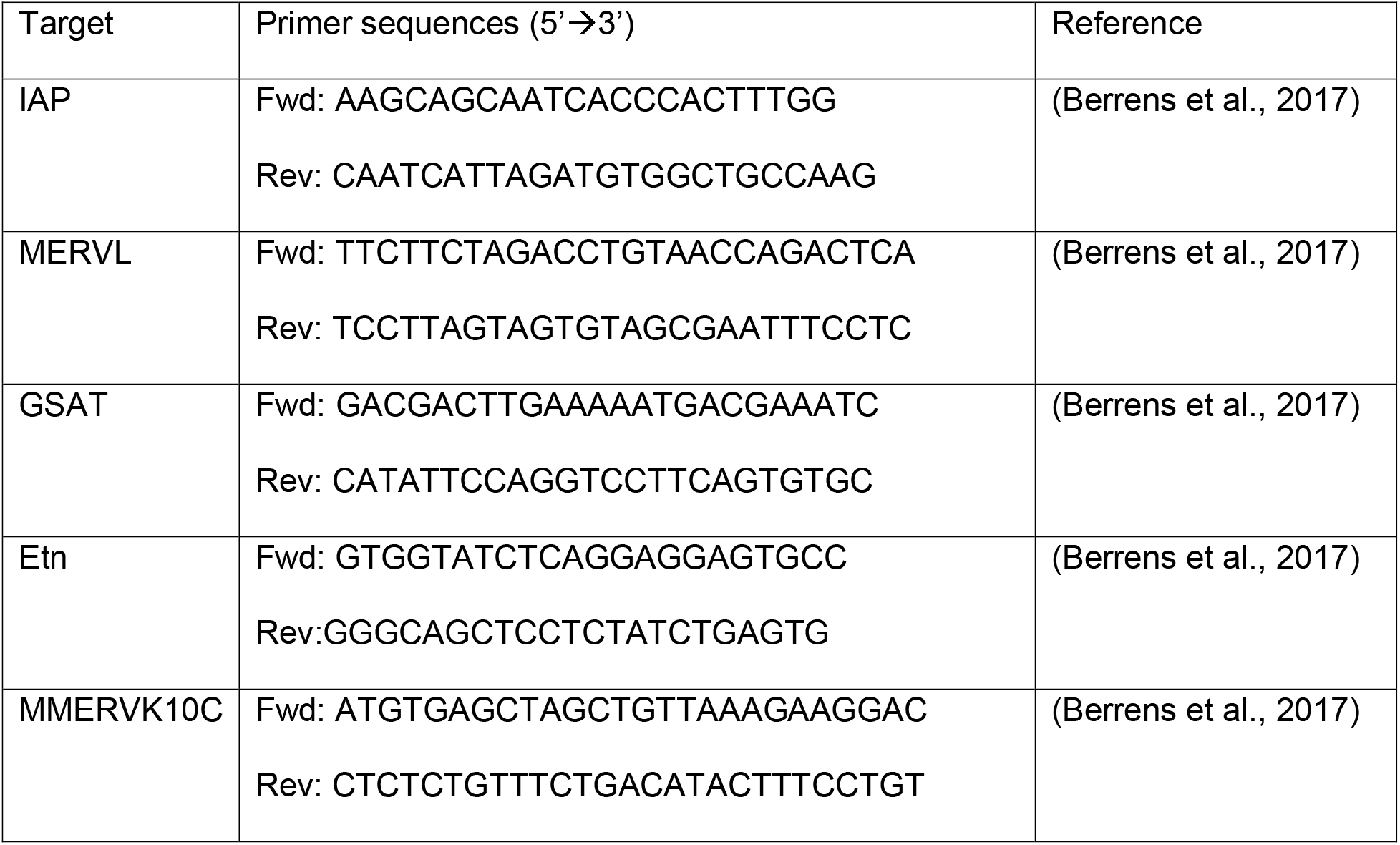
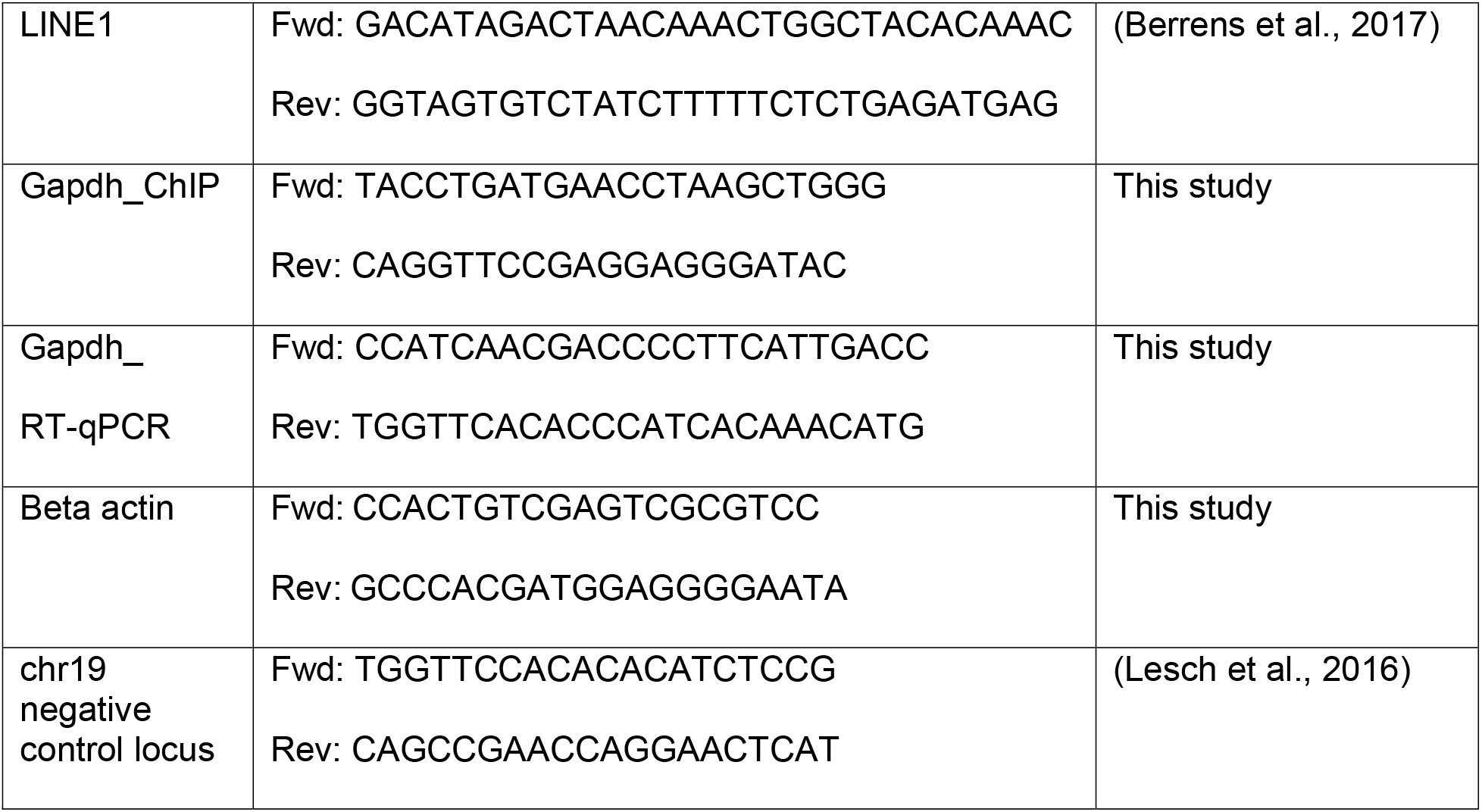

#### Generation of *Dot1L* knockout mESC lines

*Dot1L* knockout (KO) mESCs were generated using CRISPR/Cas9. Briefly, paired gRNAs were designed using E-CRISP (Heigwer et al., 2014) to target exon 5 of *Dot1L*, and cloned into pLenti-CRISPR-V2-Puro (Addgene # 52961) using the BsmBI restriction site as described previously (Shalem et al., 2014). gRNA sequences are as follows: Dot1l-01_Fwd: caccgGGTCTCCCCATACACCTCAG, Dot1l-01_Rev: aaacCTGAGGTGTATGGGGAGACCc; Dot1l-02_Fwd: caccgCCCAGATGATTGATGAGATC, Dot1l-02_Rev: aaacGATCTCATCAATCATCTGGGc. v6.5 mouse ESCs were seeded on gelatinized (0.2% gelatin) plates and transfected with gRNA vectors using the Fugene transfection reagent (BioRad). 14-18 hours after transfection, media was changed and puromycin-resistant feeder cells (Thermo Fisher #A34959) were added. 4-6h after adding feeders (∼24h after transfection), puromycin (2ug/ml) was added and transfected mESCs were selected in puromycin for 48 hours. After 7 days of selection, individual colonies were picked into 96 well plates and allowed to regrow for another 2-3 days. Cells were lysed overnight at 37C in lysis buffer (10 mM Tris-HCl, pH 7.5, 10 mM EDTA, 10 mM NaCl, 0.5% sarcosyl, 1 mg/mL proteinase K) and genomic DNA was isolated. The targeted region (exon 5) of Dot1L was PCR amplified and Sanger sequenced to confirm the deletion.

#### Generation of *Smarca5* knockdown mESC lines

Lentiviral vectors encoding either a non-targeting shRNA (TLNSU4440) or two specific shRNAs directed against mouse *Smarca5* (ULTRA3453660 and ULTRA3454387, transOMIC technologies, Cat#TLMSU14222), were co-transfected with VSV-G and psPAX2 (Addgene #12260) plasmids into HEK293T cells (RRID:CVLC_0063) using Lipofectamine 3000 transfection reagent (Invitrogen, #L3000008) and incubated at 37 °C, 5% CO_2_ to generate lentiviral particles. After 48 h, viral particles were harvested from the culture supernatant by filtering through a 0.45 μm syringe filter unit. Viral particles harboring either non-targeting control or *Smarca5* directed shRNA were used to infect v6.5 mESCs following treatment with polybrene (8 μg/mL, Millipore Sigma, # TR-1003-G) for 2 h. After 48 h, transduced cells were selected with 2 μg/mL puromycin (Gibco, #A1113803) with daily media changes. Once cells reached optimal growth and formed clearly distinguishable colonies, individual colonies were picked and expanded in a 96 well plate. All colonies were maintained in complete mESC media as described above. Selected clones were expanded into a 24 well plate and knockdown of SMARCA5 was confirmed by immunoblot analysis.

#### Metaphase spreads

V6.5 mESCs were plated on gelatinized 6-well dishes and allowed to grow for 1-2 days. 1 hour prior to addition of Colcemid (Karyomax, Gibco #15210-040), cells were fed with fresh ESC media. Cells were treated with Colcemid to a final concentration 1ug/ml and allowed to grow at 37C for 1h. After rinsing with PBS, cells were trypsinized and pelleted down at 1000rpm for 5 minutes. The supernatant was aspirated, leaving about 200ul media in the tube. The cell pellet was resuspended by gently tapping the bottom of the tube, and 5ml of ice cold 0.56% KCl solution was added slowly and mixed by inverting the tube once. Cells were incubated at room temperature for 6 min and then spun at 1000rpm for 4 min. The supernatant was aspirated and the pellet was resuspended in the remaining drops of liquid by gently tapping the bottom of the tube. Cells were fixed with 5ml of fixative (methanol: glacial acetic acid, ratio 3:1). Fixative was added dropwise with continuous mixing. After 5 minutes cells were centrifuged at 1000rpm for 4 min. Finally, cells were resuspended in 500ul of remaining fixative and released one drop at a time onto an alcohol cleaned slide with a 20ul pipettor by holding the slide slightly tilted over a jar of water. Slides were air dried for 2 hours at room temperature. After adding 1 or 2 drops of antifade DAPI mounting media (Invitrogen), slides were coverslipped and imaged on LSM 980 confocal microscope (Zeiss) as mentioned above using ZEN acquisition software.

#### Mouse embryo collection

B6D2F1 strain female mice (age 6–8 weeks, Jackson Labs) were superovulated by serial Pregnant Mare Serum Gonadotropin (5 IU per mouse, Prospec Protein Specialists) and human chorionic gonadotropin (5 IU, MilliporeSigma) injections 47 h apart. The following day, MII stage oocytes were isolated in M2 media supplemented with hyaluronidase (MilliporeSigma) and stored in 25 μl drops of pre-gassed KSOM with half-strength concentration of amino acids (MilliporeSigma) under mineral oil (Irvine Scientific). Zygotes were generated by piezo-actuated intracytoplasmic sperm injection (ICSI; see (Grosswendt et al., 2020) using thawed CAST/EiJ strain sperm in batches of 30–50 oocytes and standard micromanipulation equipment, including a Hamilton Thorne XY Infrared laser, Eppendorf Transferman NK2 micromanipulators, and a Zeiss Axio Observer inverted microscope. Alternatively, superovulated B6D2F1 females were crossed with B6D2F1 males and zygotes isolated the following morning.

For DOT1L inhibitor treatment experiments, 10 μM SGC0946 in KSOM was plated the night before and isolated oocytes were deposited directly into these drops prior to ICSI, after which they were returned to new drug containing drops and washed several times to serially dilute the M2 media. Then, progression to set morphological stages was conducted by visual inspection at 24 hpf (2-cell), 48 hpf (4-cell), 56 hpf (8-cell), 72 hpf (early morulae), 84 hpf (late morulae, early blastocyst) and 96 hpf (late blastocyst).

Embryos were prepared for fixing in 4% paraformaldehyde by removal of the zona pellucida using Acid Tyrode’s Solution (MilliporeSigma) and directly plating onto Poly-L-lysine coated dishes (MatTek) in PBS.

#### Immunofluorescence staining and imaging of mESCs

Wild-type and *Dot1L* KO ESCs were plated on fibronectin (10ug/ml) coated 35mm coverslip dishes (MatTek corp. #P35GC-1.5-14-C.S) for overnight growth. Cells were fixed in 4% paraformaldehyde for 15 minutes, washed with DPBS and permeabilized with 0.2% Triton X-100 in PBS for 10 minutes. Cells were then blocked in blocking buffer (5% BSA, 0.1% Triton X-100 or 4% FBS, 0.1% Triton X-100) for 1 hour at room temperature (RT) and then incubated overnight at 4°C with primary antibodies diluted in blocking buffer. The dishes were washed with 0.1% PBST and stained with fluorophore-conjugated secondary antibodies: AlexaFluor 488-conjugated goat anti-rabbit-IgG (Molecular Probes), AlexaFluor 568-conjugated goat anti-rabbit-IgG (Molecular Probes), AlexaFluor 568-conjugated goat anti-mouse-IgG (Molecular Probes). All secondary antibodies were used at 1:700 dilution in blocking buffer and incubated at RT for 1h. Finally, the slides were mounted in antifade mounting medium containing DAPI (Invitrogen). Images were acquired with LSM 710 or LSM 980 confocal microscope (Zeiss) equipped with 405, 488, 555/561 nm and two-photon lasers, and fitted with a 63×1.4 NA objective using ZEN acquisition software.

#### Immunofluorescence staining and imaging of embryos

Embryos were grown in vitro from 4 to 96 hours with or without treatment of DOT1L inhibitor SGC0946. Embryos were collected at the appropriate time point and fixed with 4% paraformaldehyde for 20 min at 4C. After 3 washes with PBS-1%BSA for 5 minutes each, embryos were permeabilized with 0.2% Triton X-100 for 20 min at room temperature and blocked with blocking solution (5% BSA, 0.15% Triton X-100) for 1 h at room temperature. Embryos were washed 3 times with wash buffer (0.1% PBS-Triton X-100) and incubated with primary antibodies overnight at 4C in blocking solution. The next day, embryos were washed with wash buffer and stained with fluorophore-conjugated secondary antibodies: AlexaFluor 488-conjugated goat anti-mouse-IgG (Molecular Probes) or Alexa Fluor 568-conjugated goat anti-mouse-IgG (Molecular Probes). All secondary antibodies were used at 1:700 dilution in blocking buffer and incubated at RT for 1 h. Finally, the slides were mounted in antifade mounting medium containing DAPI (Invitrogen). Images were acquired with LSM 710 or LSM 980 confocal microscope (Zeiss) equipped with 405, 488, 555/561 nm and two-photon lasers, and fitted with a 63×1.4 NA objective using ZEN acquisition software.

#### Real-time quantitative PCR

Reverse transcription of 1µg of total RNA was performed with oligo dT primers and iScript reverse transcriptase (Bio-Rad #1708896) in a total volume of 20µl according to the manufacturer’s instructions. Reaction mixtures were incubated in a thermocycler at 25°C for 5 min then 42°C for 60 min before stopping the reaction at 95°C for 1 min. 1 µl of undiluted cDNA was used for a 20ul reaction volume consisting of 4µl of 10µM forward plus reverse primer mix, 10µl of Power SYBR Green PCR Master Mix (Applied Biosystems #4367659) and 5µl nuclease-free water. Primer sequences used for qPCR are listed in the ‘Primers’ subsection above. Reactions for each target gene were performed in duplicate in a 96 well plate loaded into an Applied Biosystems QuantStudio 3 Real-Time PCR System. Standard cycling conditions were used: Hold stage (x1): 50°C for 2 min, 95°C for 10 min; PCR stage (x40): 95°C for 15 sec, 60 °C for 1 min. Melt curve stage conditions were: 95°C for 15 secs, 60°C for 1 min, 95°C for 15 secs. Relative fold change in transcript abundance was calculated using the delta-delta Ct method by normalizing target gene expression levels to either *Gapdh* or *Actb*.

#### Western blot analysis

Cells were lysed in RIPA buffer (50 mM Tris-HCl pH 7.4, 150 mM NaCl, 1% Triton X-100, 0.5% sodium deoxycholate, 0.1% SDS, 1 mM EDTA, 10 mM NaF, 1 mM PMSF) containing fresh cOmplete EDTA-free protease inhibitor cocktail (Roche). Cell extracts were subjected to sonication using a Bioruptor bath sonicator (Diagenode) and centrifuged at 20,000xg for 15 min at 4°C and supernatants were immediately mixed with 4x SDS sample buffer to a final dilution of 1x. The samples were heated at 95°C for 5 min, resolved on a Mini-PROTEAN TGX gel (Bio-Rad, 456-8093) by SDS-PAGE for 2 hour at 80V and transferred onto PVDF membranes (GE Healthcare) in 20 mM Tris-HCl (pH 8.0), 150 mM glycine, 20% methanol. Following transfer, membranes were blocked with 5% skim (non-fat) milk for 1 h at RT, followed by incubation with the primary antibody overnight at 4° C. After washing three times with TBST buffer (20 mM Tris-HCl pH 7.6, 150 mM NaCl and 0.1% Tween 20), membranes were incubated at RT for 1 h with HRP-conjugated secondary antibodies as follows: goat anti-rabbit IgG conjugated to HRP (Jackson Immuno Research, 111-035-003, 1:20,000) for H3K79me1, H3K79me2, H3K79me3, SMARCA5, DOT1L; and goat anti-mouse IgG conjugated to HRP (Jackson Immuno Research, 111-035-003, 1:20,000) for CHD4, HMGA1 and GAPDH. After washing the membrane three times with TBST, specific protein bands were detected using a SuperSignal West Pico PLUS Chemiluminescent Substrate (ThermoFisher #34577). Chemiluminescence was detected using the FluorChem E (Protein Simple) documentation system. Densitometry analysis of bands was performed by using ImageJ software (Fiji) or the multiplex band analysis tool in AlphaView software (Protein Simple). The band intensity of the protein of interest was normalized to the loading control.

#### Co-immunoprecipitation of endogenous DOT1L

Wild type and *Dot1L* KO mESCs were grown in 10cm dishes. To harvest, cells were washed with ice cold PBS twice and lysed in ice-cold lysis buffer (25mM Tris.HCl pH 7.4, 150mM NaCl, 1mM EDTA, 1% NP40, 0.5% Triton X-100, 5% glycerol, 1mM PMSF, 2mM NaF, 1X Protease inhibitor cocktail (Roche)) for 5 minutes on ice. Cells were harvested by scraping, and lysate was incubated at 4C for 30 min with constant agitation. The lysate was centrifuged at 14,000xg for 10 min at 4C and the supernatant was transferred to a fresh 1.5 ml tube. Preclearing was performed for 1h at 4C by adding 20ul of Protein G Dynabeads to the lysate. After a brief spin, supernatant was collected on a magnetic stand. 10% of the lysate was set aside as input. The remaining lysate was mixed with 10ug of primary antibody and incubated overnight at 4C with end-over-end mixing. 20ul of fresh beads were added to the lysate and incubated at 4C for 2-4h with end-over-end rotation. Beads were collected on a magnetic stand and washed 3 times in wash buffer (10mM Tris.HCl pH 7.4, 1mM EDTA, 1mM EGTA pH 8.0, 150mM NaCl, 1% Triton X-100, 0.2mM sodium orthovanadate, Protease inhibitor cocktail). Finally, the antigen-antibody complex was eluted by heating the beads in 4x SDS buffer at 95C for 10 min. Co-immunoprecipitated proteins were analyzed by SDS-PAGE.

#### Co-immunoprecipitation of HA-tagged DOT1L for mass spectrometry

*Dot1L* KO mESCs were transfected with MSCB-hDot1Lwt plasmid (Addgene #74173) expressing HA-tagged human DOT1L. HA-tagged DOT1L was immunoprecipitated from a confluent 10cm dish using the Pierce HA-Tag Magnetic IP/Co-IP Kit (Pierce™ HA-Tag Magnetic IP/Co-IP Kit, Cat # 88838) following the manufacturer’s instructions with some modifications. A corresponding untransfected dish was kept as a negative control. Briefly, transfected and untransfected *Dot1L* KO ESCs were washed with ice-cold PBS twice and lysed in ice-cold lysis buffer (25mM Tris.HCl pH 7.4, 150mM NaCl, 1mM EDTA, 1% NP40, 0.5% Triton X-100, 5% glycerol, 1mM PMSF, 2mM NaF, 1X Protease inhibitor cocktail (Roche)) for 5 minutes on ice.

Cells were harvested by scraping and lysate was incubated at 4C for 30 min with constant agitation. Lysate was centrifuged at 14,000xg for 10 min at 4C, and the supernatant was transferred to a fresh 1.5 ml tube. Preclearing was performed for 1h at 4C by adding 20ul of Protein G Dynabeads (Thermo Scientific #10007D) to the lysate. After a brief spin, supernatant was collected on a magnetic stand. 30ul (0.30mg) of Pierce Anti-HA magnetic beads was added to a fresh 1.5mL microcentrifuge tube and washed with lysis buffer several times. Lysate containing the HA-tagged protein was added to the pre-washed anti-HA magnetic beads and incubated at 4C for 8h with constant mixing. After 2 washes with lysis buffer and 1 wash with ultrapure water, beads were collected and eluted with elution buffer according to the manufacturer’s instructions. Eluted sample was mixed with 5x reducing sample buffer, heated at 95C for 10 min, and anaylzed by SDS-PAGE. For mass spec analysis, immunoprecipitated proteins were resolved for up to 1cm on an SDS-PAGE gel. The gel was fixed in 30% ethanol and 10% acetic acid in water. The gel piece was excised and submitted to the Keck MS & Proteomics Resource at Yale University for analysis.

#### Mass spectrometry sample processing and analysis

Excised 1D SDS polyacrylamide gel band/plug corresponding to co-IP eluted proteins were subjected to *in situ* enzymatic digestion. The gels are washed with 250μl 50% acetonitrile/50% water for 5 minutes followed by 250μl of 50mM ammonium bicarbonate/50% acetonitrile/50% water for 30 minutes. One final wash is done using 10mM ammonium bicarbonate/50% acetonitrile/50% water for 30 minutes. After washing, the gel plugs are dried in a Speedvac and rehydrated with 0.1μg of either modified trypsin (Promega), per (approximately) 15mm^3^ of gel in 15 μl 10mM ammonium bicarbonate. Samples are digested at 37 °C for 16 hours (overnight).

Digest is then centrifuge at supernatant/solution is transferred to injection vial to be injected onto a Waters NanoACQUITY UPLC coupled to a Q-Exactive Plus mass spectrometer. The Waters nanoACQUITY UPLC system uses a Waters Symmetry® C18 180µm x 20mm trap column and a 1.7 µm, 75 µm x 250 mm nanoAcquity™ UPLC™ column (35°C) for peptide separation.

Trapping is done at 5µl/min, 99% Buffer A (100% water, 0.1% formic acid) for 3 min. Peptide separation is performed at 300 nl/min with Buffer A: 100% water, 0.1% formic acid and Buffer B: 100% CH_3_CN, 0.075% formic acid. For the MS and MS/MS Q-Exactive Plus data collection, High-energy Collisional Dissociation (HCD) MS/MS spectra filtered by dynamic exclusion was acquired over a 3 second duty cycle for charge states 2-8 with m/z isolation window of 1.6. All MS (Profile) and MS/MS (centroid) peaks were detected in the Orbitrap. Trapping was carried out for 3 min at 5 µl/min in 97% Buffer A (0.1% FA in water) and 3% Buffer B [(0.075% FA in acetonitrile (ACN)] prior to eluting with linear gradients that will reach 5% B at 1 min, 255% B at 90 min, and 50% B at 110 min, and 90% B at 115 min for 5 min; then dropdown to 3% B at 125 min for 5 min.

LC MS/MS data were analyzed utilizing Proteome Discoverer 2.4 (Thermo Fisher Scientific) with Mascot search engine (v. 2.7 Matrix Science LLC.). Resulting PD analyses were imported into Scaffold (v. 4.0, Proteome Software) for further interrogation. Positive protein identification were based on hits with two or more unique peptides per protein, and peptides were considered significant if the Mascot Score is better than the 95% confidence level.

#### Chromatin immunoprecipitation (ChIP)

Cells were cross-linked with 1% formaldehyde at room temperature for 10 min. Formaldehyde was quenched with 0.1375M glycine at room temperature for 10 minutes. Fixed cells were spun down at 1500rpm for 5 minutes at 4°C. Cells were washed with cold PBS twice and resuspended in 50ul ChIP lysis buffer (1% SDS, 10mM EDTA, 50mM Tri-HCl at pH 8.1) and frozen in -80C. Antibody-bound Dynabeads (Thermo Fisher #10007D) were prepared by mixing 10 μ l block solution (0.5% BSA in PBS). Beads were washed twice with 150 μl of block solution and resuspended in 30ul block solution. 1ug antibody per million cells for H3K79me2, H3K79me3, HP1β, or RNA PolIIS2P antibodies was added to each aliquot of beads and incubated for 8 hours rotating at 4°C. Between 2-4×10^6^ cells were used for ChIP-seq and ChIP-qPCR. Frozen cells were thawed on ice and ChIP dilution buffer (0.01% SDS, 1.1% Triton X-100, 1.2mM EDTA, 167mM NaCl, 16.7mM Tris-HCl at pH 8.1) was added to reach a total volume of 150 μl. Cells were sonicated at 4°C for 30 cycles (30 seconds on/off) using a Bioruptor bath sonicator (Diagenode). Aliquots of the same samples were pooled and spun down at 12,000xg for 5 min. The supernatant was moved to a new 1.5 ml Eppendorf tube and 600 μ l protease inhibitor cocktail (Roche #11836153001) were added. 50 μl of each sample was set aside as input before an aliquot of primary antibody-bound Dynabeads was added to the lysate and incubated overnight rotating at 4°C. After overnight incubation, beads were washed twice with low-salt immune complex wash buffer (0.1% SDS, 1% Triton X-100, 2mM EDTA, 150mM NaCl, and 20mM Tris-HCL at pH 8.1), twice with LiCl wash buffer (0.25M LiCl, 1% NP40, 1% deoxycholate, 1mM EDTA, and 10mM Tris-HCl at pH 8.1), and twice with TE (1 mM EDTA and 10mM Tris-HCl at pH 8.0). Bound DNA was eluted twice with 125 μl elution buffer (0.2% SDS, 0.1M NaHCO3, and 5mM DTT in TE) at 65°C and crosslink reversal was performed by incubating at 65°C for 8-15h. ChIP and input samples were incubated for 3 hours at 37°C with 0.2 mg/ml RNAse A (Millipore 70856-3), and 4-10 hours at 55°C with 0.1 mg/ml Proteinase K (NEB P8107S). Sample DNA was prepared using a Zymo ChIP DNA Clean & Concentrator kit (Zymo Research #D5201) according to the manufacturer’s instructions. Columns were washed twice with 200 μ l Elution Buffer into fresh Eppendorf tubes, then re-eluted with the same eluate to enhance the yield.

#### RNA isolation and sequencing library preparation

For mESCs, the cell pellet was resuspended in 1 mL of TRIzol and cells were homogenized by drawing up and down in a 21G needle with 3 ml syringe. 200ul chloroform was added, then samples were vortexed and incubated briefly at room temperature and centrifuged at 12000xg, 15 min, 4C. The aqueous phase was transferred to a gDNA eliminator column from the RNEasy Micro Plus kit (Qiagen), and total RNA was isolated according to manufacturer’s instructions. For embryos, two replicates of 30-35 mouse embryos at 56 hours post fertilization were each disrupted in 75 ul Buffer RLT, and total RNA was prepared using the RNeasy Micro kit (Qiagen #74004) according to the manufacturer’s instructions. RNA was eluted in 12ul RNase-free water.

#### Sequencing library preparation

DNA or RNA integrity and fragment size were confirmed on a BioAnalyzer (Agilent #G2939BA). For ChIP-seq, approximately 5-10ng of DNA was end-repaired, A-tailed, adapter-ligated, and PCR enriched (8-10 cycles) using the KAPA Hyper Library Preparation kit (KAPA Biosystems, #KK8504) according to the manufacturers’ instructions. For RNA-seq, libraries were prepared by ribosome depletion using the KAPA RNA Hyperprep Kit with RiboErase (Roche #08098140702) (mESCs) or the SMARTer Stranded Total RNA-seq kit v3 – Pico Input (Takara Bio #634485) (embryos) according to the manufacturers’ instructions. Indexed libraries were quantitated by qPCR using the KAPA Library Quantification Kit (KAPA Biosystems #KK4854). All samples were sequenced on an Illumina NovaSeq using 100bp paired-end sequencing. De-multiplexing was performed using CASAVA 1.8.2 (Illumina).

#### RNA-seq data analysis

For single-copy genes, data were aligned and assembled with kallisto (Bray et al., 2016) or STAR (Dobin et al., 2013) using default parameters and mouse genome assembly mm10. Read counts from STAR alignments were generated using HTSeq (Anders et al., 2014). Differential gene expression was called using DESeq2 (Love et al., 2014), with a cutoff p-value of < 0.05 after adjusting for multiple comparisons. For analysis of repetitive elements, data were first aligned to the mm10 genome assembly using Bowtie2 (Langmead and Salzberg, 2012) and multi-aligning reads were parsed using RepEnrich2 (Criscione et al., 2014). Differential repeat expression was called using EdgeR (Robinson et al., 2010) according to the RepEnrich2 pipeline. Plotting was done in R using the ggplot2 (Wickham, 2016) and lattice (Sarkar, 2008) packages.

#### ChIP-seq data analysis

Data were filtered for high-quality reads using the fastq_quality_filter tool from FASTX-Toolkit (Hannon, 2010) with parameters -q 20 -p 80 and aligned to the mm10 genome assembly using Bowtie2 with –end-to-end --fast parameters. Peaks were called using MACS2 with default parameters (Zhang et al., 2008). Peak intersections were evaluated using BEDTools (Quinlan and Hall, 2010). Analysis of repetitive elements was performed using RepEnrich2, and repeats were called as significantly enriched at false discovery rate < 0.05 for ChIP compared to input control. Peaks were assigned to genes using GREAT (McLean et al., 2010). Metagene profiles were generated with the plotProfile function within the deepTools software (Ramírez et al., 2016).

#### Gene Ontology analysis

Analysis of functional enrichments for RNA-seq, ChIP-seq, and proteomics data was performed using the Molecular Signatures Database tool (MSigDB) (Liberzon et al., 2011; Subramanian et al., 2005) to search the Gene Ontology Biological Function database (Ashburner et al., 2000; Gene Ontology Consortium, 2021).

### QUANTIFICATION AND STATISTICAL ANALYSIS

Assays were done in triplicate (n=3) unless stated otherwise in the figure legend, and the meaning of n is described in each figure legend. Error bars represent standard deviation or standard error of the mean as specified in the relevant figure legend. Statistical comparisons were performed using unpaired Student’s t-test for continuous variables and using Fisher’s Exact test for proportions. When multiple comparisons were performed, p-values were corrected as described in the figure legends or relevant Methods sections. Differences were considered statistically significant at p<0.05 (for single comparisons) or false discovery rate (FDR) <0.05 (for multiple comparisons). Statistics for genomics and proteomics data analysis are described in the relevant Methods sections above.

## Supplemental Information titles and legends

**Table S1.** Summary of high-throughput sequencing libraries.

**Table S2.** ChIP-seq peak coordinates, associated genes, and enriched GO terms.

**Table S3.** mESC RNA-seq differentially expressed genes and enriched GO terms.

**Table S4.** IP-mass spec interactor list and enriched GO terms.

**Table S5.** Embryo RNA-seq differentially expressed genes and enriched GO terms.

---

**Figure S1. Related to Figure 1.**
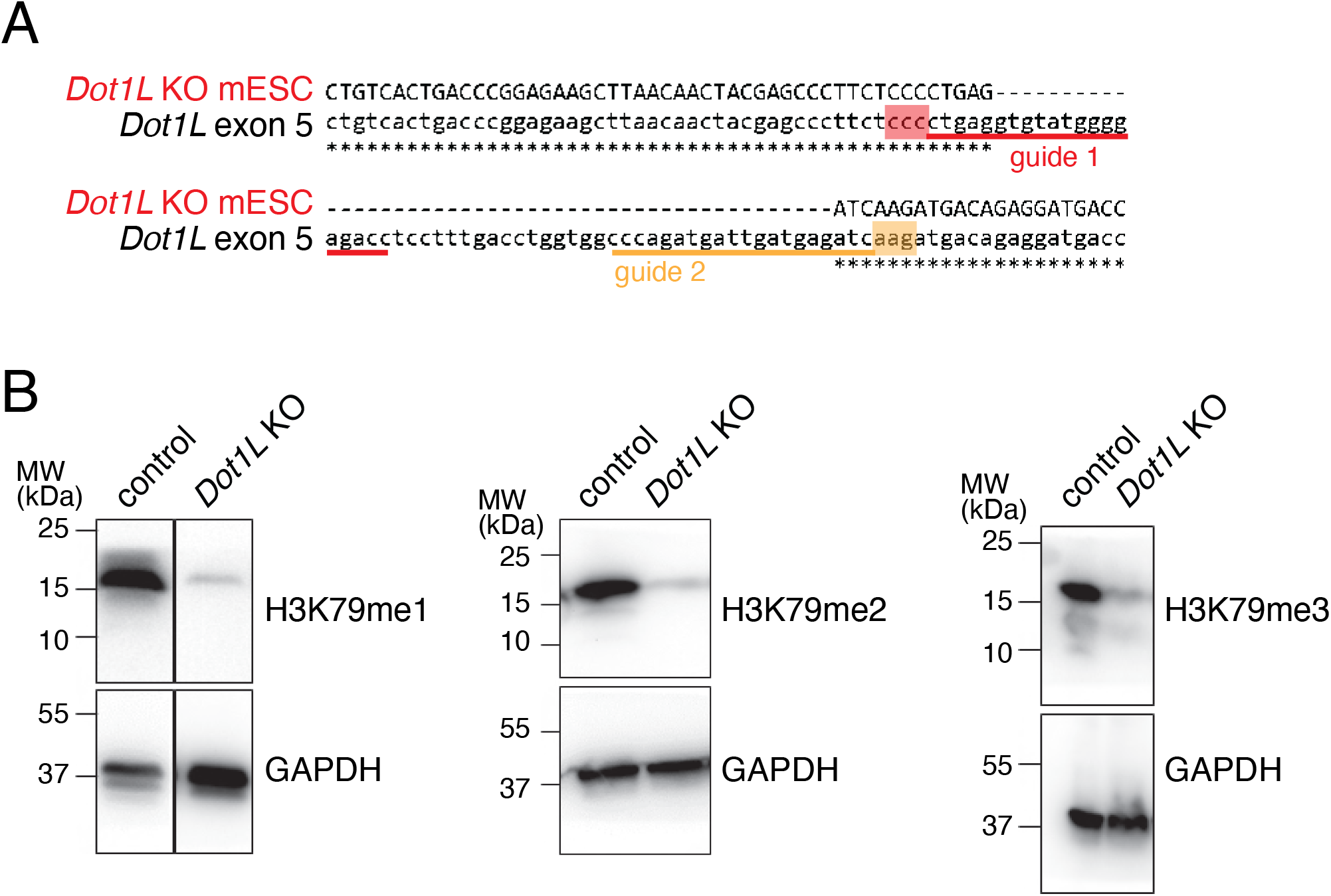
Validation of Dot1L knockout mESCs. **A,** Partial sequence of *Dot1L* exon 5 showing the sequence deleted in *Dot1L* KO cells, and the sequences (red and orange lines) and PAM sites (red and orange boxes) for the two guide RNAs used to generate the knockout. **B,** Western blots showing H3K79 methylation in *Dot1L* KO cells. GAPDH was used as a loading control. mESCs were grown on feeder cells, which are the likely source of residual bands seen in *Dot1L* KO cells.

**Figure S2. Related to Figure 1.**
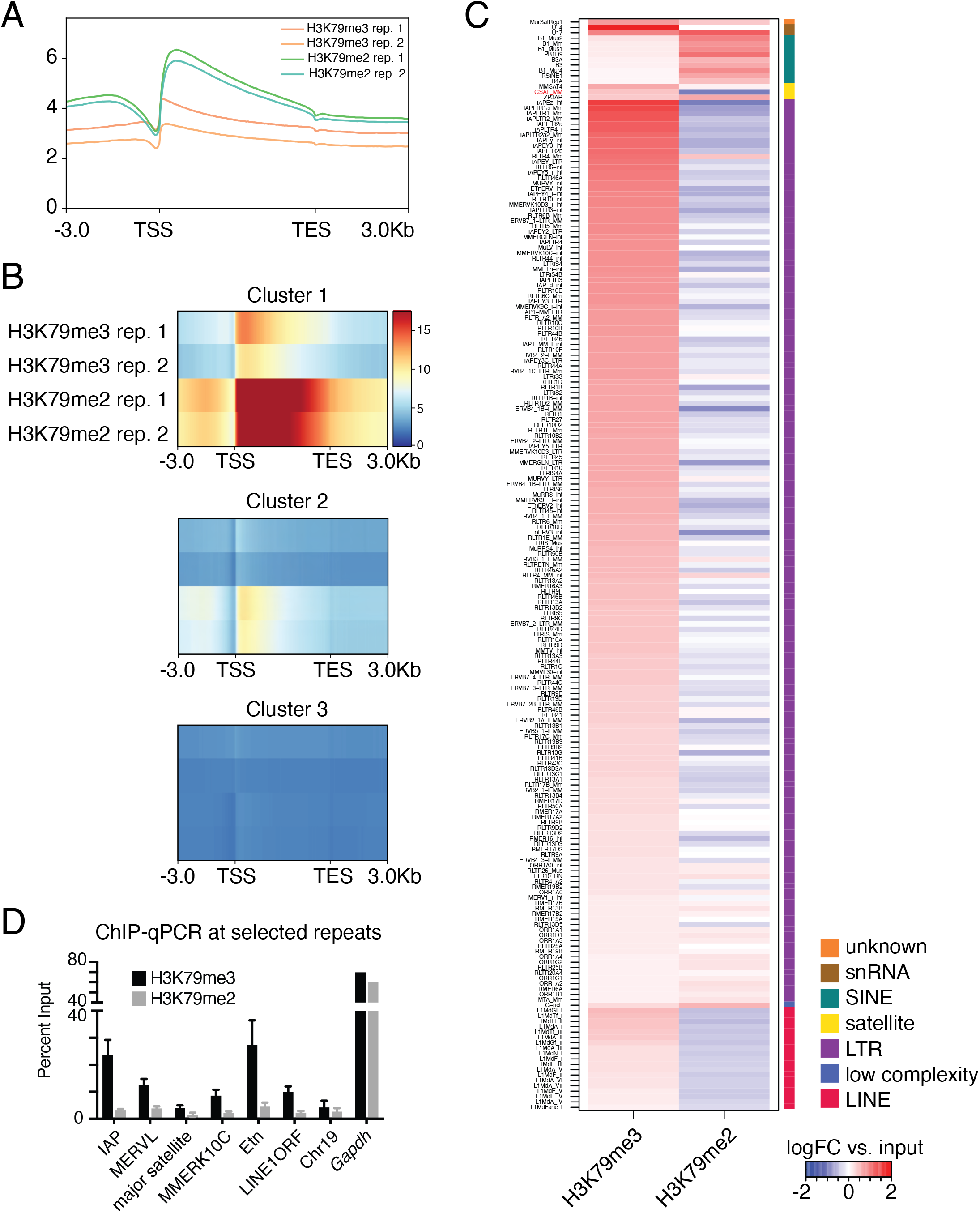
H3K79me3 is enriched at repetitive sequences in mESCs. **A,** Metagene profiles of average H3K79me2 and H3K79me3 signal at single-copy transcripts. **B,** Metagene heatmaps for H3K79me2 and H3K79me3 in three transcript clusters defined by k-means clustering. Cluster 1 includes transcripts (n=10972) positive for both H3K79me2 and H3K79me3, cluster 2 includes transcripts (n=40619) positive for H3K79me2 only, and cluster 3 includes transcripts (n=90138) negative for both. **C,** Heatmap showing enrichment of ChIP vs. input tags at repeat element annotations for H3K79me3 and H3K79me2. Elements are ordered based on RepeatMasker class and then by H3K79me3 enrichment. The major satellite repeat (GSAT_MM) is highlighted in red. **D,** ChIP-qPCR for H3K79me3 and H3K79me2 at selected repeat elements. Chr19 is a negative control locus, and Gapdh is an active single-copy locus that serves as a positive control.

**Figure S3. Related to Figure 2.**
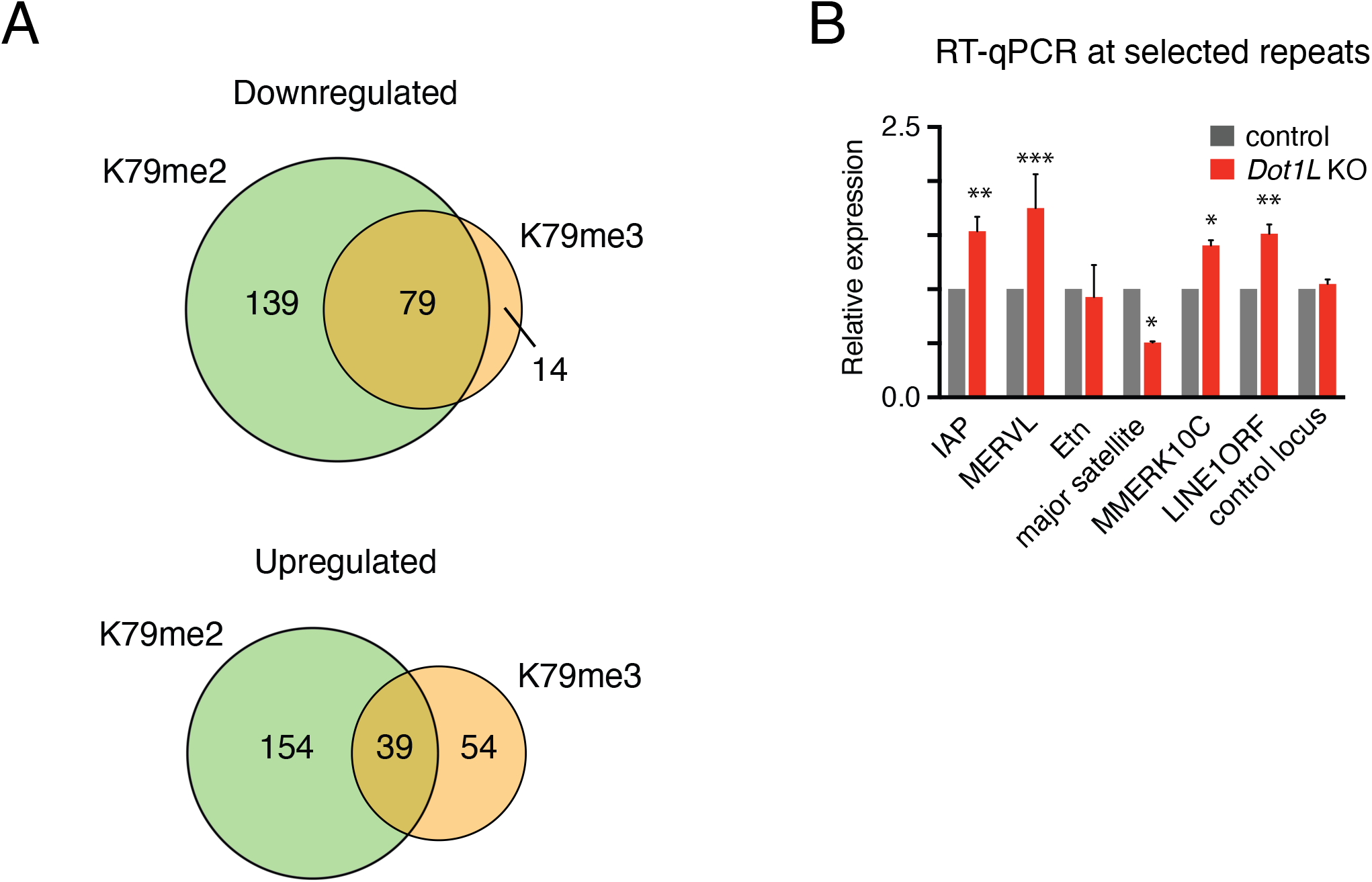
Levels of major satellite transcripts are dependent on DOT1L. **A,** Gene sets that are transcriptionally downregulated (top) or upregulated (bottom) and marked by H3K79me2, H3K79me3, or both in *Dot1L* KO mESCs. **B,** RT-qPCR for selected repeat element transcripts in wild type and *Dot1L* KO mESCs. Bars represent mean of 3 biological replicates and error bars represent ± SEM. *p<0.05, **p<0.01, ***p<0.001, unpaired Student’s t-test.

**Figure S4. Related to Figure 4.**
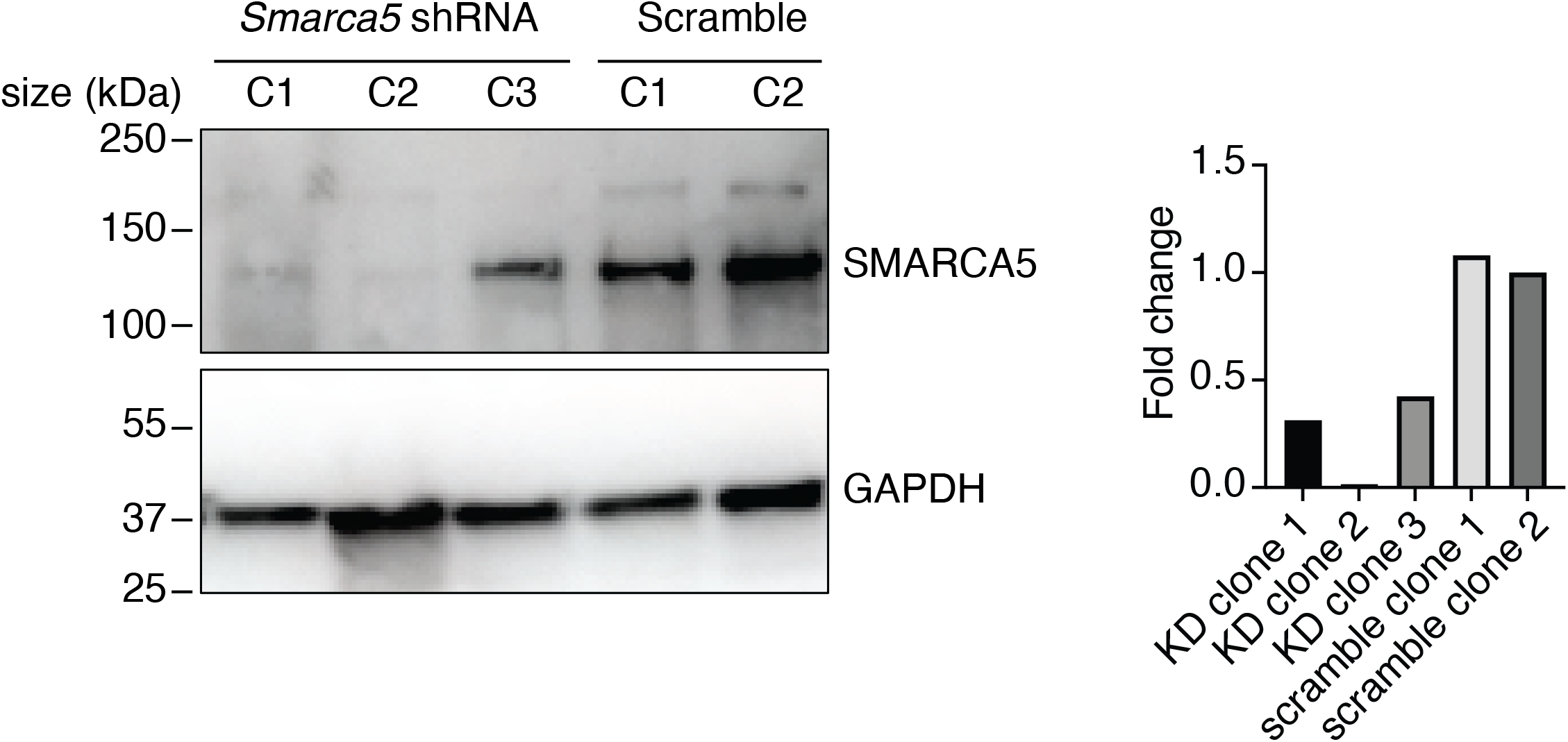
Smarca5 knockdown in mESCs. Western blot showing levels of SMARCA5 in mESC clones (C1, C2, or C3) expressing shRNA against *Smarca5* or scrambled control. At right, quantitation of SMARCA5 signal normalized to GAPDH, relative to scrambled control clone #2.

**Figure S5. Related to Figure 6.**
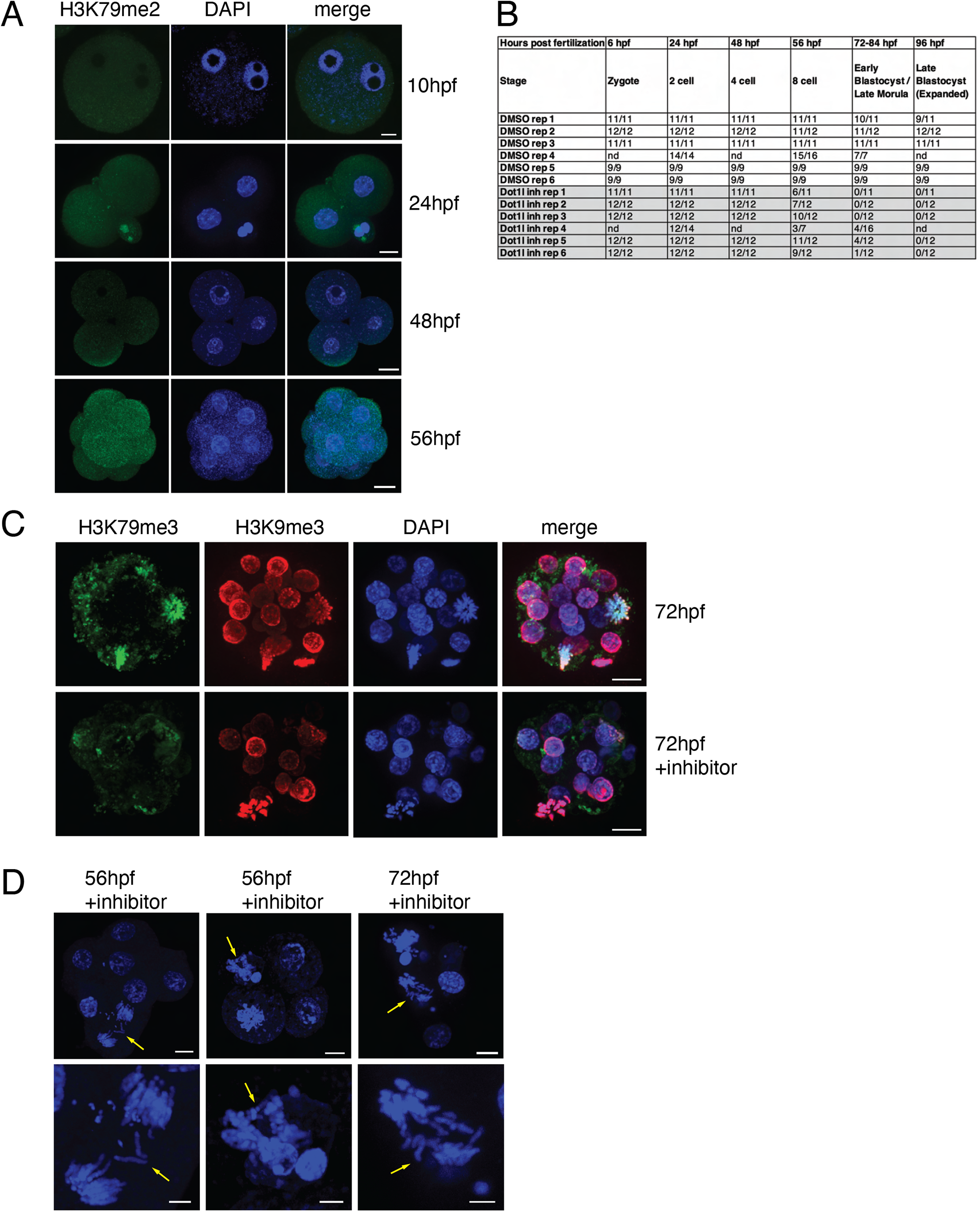
DOT1L is active in preimplantation embryos and required for embryo viability. **A,** H3K79me2 immunofluorescence in wild type embryos during preimplantation development. DNA is stained with DAPI (blue). Scale bar, 10µm. **B,** Embryo counts corresponding to the summary data displayed in Figure 6C. **C,** H3K79me3 and H3K9me3 immunofluorescence in control and DOT1L inhibitor-treated embryos at 72 hpf. DNA is stained with DAPI (blue). Scale bar, 10μm. D, DAPI-stained images of DOT1L inhibitor-treated embryos showing lagging chromosomes at anaphase (arrows). Bottom row shows increased magnification of anaphase nuclei from the image above. Scale bar, 10μm (top row), 4μm (bottom row).

## References

Abe, Y., Sako, K., Takagaki, K., Hirayama, Y., Uchida, K.S.K., Herman, J.A., DeLuca, J.G., and Hirota, T. (2016). HP1-Assisted Aurora B Kinase Activity Prevents Chromosome Segregation Errors. Developmental Cell 36, 487–497.

Aihara, T., Miyoshi, Y., Koyama, K., Suzuki, M., Takahashi, E., Monden, M., and Nakamura, Y. (1998). Cloning and mapping of SMARCA5 encoding hSNF2H, a novel human homologue of Drosophila ISWI. CGR 81, 191–193.

Almouzni, G., and Probst, A.V. (2011). Heterochromatin maintenance and establishment: Lessons from the mouse pericentromere. Nucleus 2, 332–338.

Anders, S., Pyl, P.T., and Huber, W. (2014). HTSeq-a Python framework to work with high-throughput sequencing data. Bioinformatics.

Ashburner, M., Ball, C.A., Blake, J.A., Botstein, D., Butler, H., Cherry, J.M., Davis, A.P., Dolinski, K., Dwight, S.S., Eppig, J.T., et al. (2000). Gene Ontology: tool for the unification of biology. Nature Genetics 25, 25–29.

Bernt, K.M., Zhu, N., Sinha, A.U., Vempati, S., Faber, J., Krivtsov, A.V., Feng, Z., Punt, N., Daigle, A., Bullinger, L., et al. (2011). MLL-rearranged leukemia is dependent on aberrant H3K79 methylation by DOT1L. Cancer Cell 20, 66–78.

Berrens, R.V., Andrews, S., Spensberger, D., Santos, F., Dean, W., Gould, P., Sharif, J., Olova, N., Chandra, T., Koseki, H., et al. (2017). An endosiRNA-Based Repression Mechanism Counteracts Transposon Activation during Global DNA Demethylation in Embryonic Stem Cells. Cell Stem Cell 21, 694–703.e7.

Bersani, F., Lee, E., Kharchenko, P.V., Xu, A.W., Liu, M., Xega, K., MacKenzie, O.C., Brannigan, B.W., Wittner, B.S., Jung, H., et al. (2015). Pericentromeric satellite repeat expansions through RNA-derived DNA intermediates in cancer. PNAS 112, 15148–15153.

Biscotti, M.A., Canapa, A., Forconi, M., Olmo, E., and Barucca, M. (2015). Transcription of tandemly repetitive DNA: functional roles. Chromosome Res 23, 463–477.

Bitoun, E., Oliver, P.L., and Davies, K.E. (2007). The mixed-lineage leukemia fusion partner AF4 stimulates RNA polymerase II transcriptional elongation and mediates coordinated chromatin remodeling. Hum Mol Genet 16, 92–106.

Borosha, S., Ratri, A., Housami, S.M., Rai, S., Ghosh, S., Malcom, C.A., Chakravarthi, V.P., Vivian, J.L., Fields, T.A., Rumi, M.A.K., et al. (2020). DOT1L primarily acts as a transcriptional repressor in hematopoietic progenitor cells.

Bray, N.L., Pimentel, H., Melsted, P., and Pachter, L. (2016). Near-optimal probabilistic RNA-seq quantification. Nature Biotechnology 34, 525–527.

Burton, A., and Torres-Padilla, M.-E. (2010). Epigenetic reprogramming and development: a unique heterochromatin organization in the preimplantation mouse embryo. Briefings in Functional Genomics 9, 444–454.

Burton, A., and Torres-Padilla, M.-E. (2014). Chromatin dynamics in the regulation of cell fate allocation during early embryogenesis. Nat Rev Mol Cell Biol 15, 723–735.

Camacho, J., Truong, L., Kurt, Z., Chen, Y.W., Morselli, M., Gutierrez, G., Pellegrini, M., Yang, X., and Allard, P. (2018). The Memory of Environmental Chemical Exposure in C. elegans Is Dependent on the Jumonji Demethylases jmjd-2 and jmjd-3/utx-1. Cell Reports 23, 2392–2404.

Cao, K., Ugarenko, M., Ozark, P.A., Wang, J., Marshall, S.A., Rendleman, E.J., Liang, K., Wang, L., Zou, L., Smith, E.R., et al. (2020). DOT1L-controlled cell-fate determination and transcription elongation are independent of H3K79 methylation. PNAS 117, 27365–27373.

Casanova, M., Pasternak, M., El Marjou, F., Le Baccon, P., Probst, A.V., and Almouzni, G. (2013). Heterochromatin Reorganization during Early Mouse Development Requires a Single-Stranded Noncoding Transcript. Cell Reports 4, 1156–1167.

Clapier, C.R., and Cairns, B.R. (2009). The Biology of Chromatin Remodeling Complexes. Annu. Rev. Biochem. 78, 273–304.

Criscione, S.W., Zhang, Y., Thompson, W., Sedivy, J.M., and Neretti, N. (2014). Transcriptional landscape of repetitive elements in normal and cancer human cells. BMC Genomics 15, 583.

Dobin, A., Davis, C.A., Schlesinger, F., Drenkow, J., Zaleski, C., Jha, S., Batut, P., Chaisson, M., and Gingeras, T.R. (2013). STAR: ultrafast universal RNA-seq aligner. Bioinformatics 29, 15–21.

Frescas, D., Guardavaccaro, D., Kuchay, S.M., Kato, H., Poleshko, A., Basrur, V., Elenitoba-Johnson, K.S., Katz, R.A., and Pagano, M. (2008). KDM2A represses transcription of centromeric satellite repeats and maintains the heterochromatic state. Cell Cycle 7, 3539–3547.

Fukagawa, T., Nogami, M., Yoshikawa, M., Ikeno, M., Okazaki, T., Takami, Y., Nakayama, T., and Oshimura, M. (2004). Dicer is essential for formation of the heterochromatin structure in vertebrate cells. Nat Cell Biol 6, 784–791.

Gene Ontology Consortium (2021). The Gene Ontology resource: enriching a GOld mine. Nucleic Acids Res 49, D325–D334.

Grewal, S.I.S., and Jia, S. (2007). Heterochromatin revisited. Nat Rev Genet 8, 35–46.

Grosswendt, S., Kretzmer, H., Smith, Z.D., Kumar, A.S., Hetzel, S., Wittler, L., Klages, S., Timmermann, B., Mukherji, S., and Meissner, A. (2020). Epigenetic regulator function through mouse gastrulation. Nature 584, 102–108.

Guenther, M.G., Lawton, L.N., Rozovskaia, T., Frampton, G.M., Levine, S.S., Volkert, T.L., Croce, C.M., Nakamura, T., Canaani, E., and Young, R.A. (2008). Aberrant chromatin at genes encoding stem cell regulators in human mixed-lineage leukemia. Genes Dev 22, 3403–3408.

Hahn, M., Dambacher, S., Dulev, S., Kuznetsova, A.Y., Eck, S., Wörz, S., Sadic, D., Schulte, M., Mallm, J.-P., Maiser, A., et al. (2013). Suv4-20h2 mediates chromatin compaction and is important for cohesin recruitment to heterochromatin. Genes Dev. 27, 859–872.

Hannon, G.J. (2010). FASTX-Toolkit.

Heigwer, F., Kerr, G., and Boutros, M. (2014). E-CRISP: fast CRISPR target site identification. Nat Methods 11, 122–123.

Hyun, K., Jeon, J., Park, K., and Kim, J. (2017). Writing, erasing and reading histone lysine methylations. Exp Mol Med 49, e324–e324.

Jagannathan, M., Cummings, R., and Yamashita, Y.M. (2018). A conserved function for pericentromeric satellite DNA. ELife 7.

Janssen, A., Colmenares, S.U., and Karpen, G.H. (2018). Heterochromatin: Guardian of the Genome. Annu Rev Cell Dev Biol 34, 265–288.

Jones, B., Su, H., Bhat, A., Lei, H., Bajko, J., Hevi, S., Baltus, G.A., Kadam, S., Zhai, H., Valdez, R., et al. (2008). The histone H3K79 methyltransferase Dot1L is essential for mammalian development and heterochromatin structure. PLoS Genetics 4, e1000190.

Jonkers, I., Kwak, H., and Lis, J.T. (2014). Genome-wide dynamics of Pol II elongation and its interplay with promoter proximal pausing, chromatin, and exons. ELife 3, e02407.

Kim, S.K., Jung, I., Lee, H., Kang, K., Kim, M., Jeong, K., Kwon, C.S., Han, Y.M., Kim, Y.S., Kim, D., et al. (2012). Human histone H3K79 methyltransferase DOT1L protein [corrected] binds actively transcribing RNA polymerase II to regulate gene expression. The Journal of Biological Chemistry 287, 39698–39709.

Kim, W., Choi, M., and Kim, J.-E. (2014). The histone methyltransferase Dot1/DOT1L as a critical regulator of the cell cycle. Cell Cycle 13, 726–738.

Kokavec, J., Zikmund, T., Savvulidi, F., Kulvait, V., Edelmann, W., Skoultchi, A.I., and Stopka, T. (2017). The ISWI ATPase Smarca5 (Snf2h) Is Required for Proliferation and Differentiation of Hematopoietic Stem and Progenitor Cells. Stem Cells 35, 1614–1623.

Langmead, B., and Salzberg, S.L. (2012). Fast gapped-read alignment with Bowtie 2. Nat Methods 9, 357–359.

Lesch, B.J., Silber, S.J., McCarrey, J.R., and Page, D.C. (2016). Parallel evolution of male germline epigenetic poising and somatic development in animals. Nature Genetics 48, 888–894.

Liao, J., and Szabó, P.E. (2020). Maternal DOT1L is dispensable for mouse development. Sci Rep 10, 20636.

Liberzon, A., Subramanian, A., Pinchback, R., Thorvaldsdóttir, H., Tamayo, P., and Mesirov, J.P. (2011). Molecular signatures database (MSigDB) 3.0. Bioinformatics 27, 1739–1740.

Liu, H., Kim, J.-M., and Aoki, F. (2004). Regulation of histone H3 lysine 9 methylation in oocytes and early pre-implantation embryos. Development 131, 2269–2280.

Love, M.I., Huber, W., and Anders, S. (2014). Moderated estimation of fold change and dispersion for RNA-seq data with DESeq2. Genome Biology 15, 550.

Lu, J., and Gilbert, D.M. (2007). Proliferation-dependent and cell cycle–regulated transcription of mouse pericentric heterochromatin. Journal of Cell Biology 179, 411–421.

McLean, C.Y., Bristor, D., Hiller, M., Clarke, S.L., Schaar, B.T., Lowe, C.B., Wenger, A.M., and Bejerano, G. (2010). GREAT improves functional interpretation of cis-regulatory regions. Nat Biotechnol 28, 495–501.

Mohan, M., Herz, H.M., Takahashi, Y.H., Lin, C., Lai, K.C., Zhang, Y., Washburn, M.P., Florens, L., and Shilatifard, A. (2010). Linking H3K79 trimethylation to Wnt signaling through a novel Dot1-containing complex (DotCom). Genes Dev 24, 574–589.

Müller, S., and Almouzni, G. (2017). Chromatin dynamics during the cell cycle at centromeres. Nat Rev Genet 18, 192–208.

Nair, L., Chung, H., and Basu, U. (2020). Regulation of long non-coding RNAs and genome dynamics by the RNA surveillance machinery. Nat Rev Mol Cell Biol 21, 123–136.

Ng, H.H., Feng, Q., Wang, H., Erdjument-Bromage, H., Tempst, P., Zhang, Y., and Struhl, K. (2002). Lysine methylation within the globular domain of histone H3 by Dot1 is important for telomeric silencing and Sir protein association. Genes Dev 16, 1518–1527.

Novo, C.L., Tang, C., Ahmed, K., Djuric, U., Fussner, E., Mullin, N.P., Morgan, N.P., Hayre, J., Sienerth, A.R., Elderkin, S., et al. (2016). The pluripotency factor Nanog regulates pericentromeric heterochromatin organization in mouse embryonic stem cells. Genes Dev. 30, 1101–1115.

Ooga, M., Inoue, A., Kageyama, S., Akiyama, T., Nagata, M., and Aoki, F. (2008). Changes in H3K79 Methylation During Preimplantation Development in Mice. Biology of Reproduction 78, 413–424.

Peters, A.H.F.M., O’Carroll, D., Scherthan, H., Mechtler, K., Sauer, S., Schöfer, C., Weipoltshammer, K., Pagani, M., Lachner, M., Kohlmaier, A., et al. (2001). Loss of the Suv39h Histone Methyltransferases Impairs Mammalian Heterochromatin and Genome Stability. Cell 107, 323–337.

Probst, A.V., Okamoto, I., Casanova, M., El Marjou, F., Le Baccon, P., and Almouzni, G. (2010). A strand-specific burst in transcription of pericentric satellites is required for chromocenter formation and early mouse development. Developmental Cell 19, 625–638.

Quinlan, A.R., and Hall, I.M. (2010). BEDTools: a flexible suite of utilities for comparing genomic features. Bioinformatics 26, 841–842.

Ramírez, F., Ryan, D.P., Grüning, B., Bhardwaj, V., Kilpert, F., Richter, A.S., Heyne, S., Dündar, F., and Manke, T. (2016). deepTools2: a next generation web server for deep-sequencing data analysis. Nucleic Acids Research 44, W160–W165.

Robinson, M.D., McCarthy, D.J., and Smyth, G.K. (2010). edgeR: a Bioconductor package for differential expression analysis of digital gene expression data. Bioinformatics 26, 139–140.

Rudert, F., Bronner, S., Garnier, J.M., and Dollé, P. (1995). Transcripts from opposite strands of gamma satellite DNA are differentially expressed during mouse development. Mamm Genome 6, 76–83.

Saksouk, N., Simboeck, E., and Dejardin, J. (2015). Constitutive heterochromatin formation and transcription in mammals. Epigenetics & Chromatin 8, 3.

Sarkar, D. (2008). Lattice: Multivariate Data Visualization with R (New York: Springer).

Shalem, O., Sanjana, N.E., Hartenian, E., Shi, X., Scott, D.A., Mikkelsen, T.S., Heckl, D., Ebert, B.L., Root, D.E., Doench, J.G., et al. (2014). Genome-Scale CRISPR-Cas9 Knockout Screening in Human Cells. Science 343, 84–87.

Shestakova, E.A., Mansuroglu, Z., Mokrani, H., Ghinea, N., and Bonnefoy, E. (2004). Transcription factor YY1 associates with pericentromeric γ satellite DNA in cycling but not in quiescent (G0) cells. Nucleic Acids Research 32, 4390–4399.

Shirai, A., Kawaguchi, T., Shimojo, H., Muramatsu, D., Ishida-Yonetani, M., Nishimura, Y., Kimura, H., Nakayama, J., and Shinkai, Y. (2017). Impact of nucleic acid and methylated H3K9 binding activities of Suv39h1 on its heterochromatin assembly. ELife 6, e25317.

Steger, D.J., Lefterova, M.I., Ying, L., Stonestrom, A.J., Schupp, M., Zhuo, D., Vakoc, A.L., Kim, J.E., Chen, J., Lazar, M.A., et al. (2008). DOT1L/KMT4 recruitment and H3K79 methylation are ubiquitously coupled with gene transcription in mammalian cells. Mol Cell Biol 28, 2825–2839.

Stopka, T., and Skoultchi, A.I. (2003). The ISWI ATPase Snf2h is required for early mouse development. PNAS 100, 14097–14102.

Stopka, T., Blafkova, J., Zakova, D., Fuchs, O., Cmejla, R., Necas, E., Jelinek, J., and Zivny, J. (2000). Cloning and expression of murine hematopoietic specific chromatin remodeling gene SMARCA5. Experimental Hematology 28, 119.

Subramanian, A., Tamayo, P., Mootha, V.K., Mukherjee, S., Ebert, B.L., Gillette, M.A., Paulovich, A., Pomeroy, S.L., Golub, T.R., Lander, E.S., et al. (2005). Gene set enrichment analysis: a knowledge-based approach for interpreting genome-wide expression profiles. Proceedings of the National Academy of Sciences of the United States of America 102, 15545–15550.

Ting, D.T., Lipson, D., Paul, S., Brannigan, B.W., Akhavanfard, S., Coffman, E.J., Contino, G., Deshpande, V., Iafrate, A.J., Letovsky, S., et al. (2011). Aberrant Overexpression of Satellite Repeats in Pancreatic and Other Epithelial Cancers. Science 331, 593–596.

Vargova, J., Vargova, K., Skoultchi, A.I., and Stopka, T. (2009). Nuclear localization of ISWI ATPase Smarca5 (Snf2h) in mouse. Frontiers in Bioscience 1, 553–559.

Veloso, A., Kirkconnell, K.S., Magnuson, B., Biewen, B., Paulsen, M.T., Wilson, T.E., and Ljungman, M. (2014). Rate of elongation by RNA polymerase II is associated with specific gene features and epigenetic modifications. Genome Res. 24, 896–905.

Wickham, H. (2016). ggplot2: Elegant Graphics for Data Analysis (New York: Springer-Verlag).

Wood, K., Tellier, M., and Murphy, S. (2018). DOT1L and H3K79 Methylation in Transcription and Genomic Stability. Biomolecules 8.

Wu, A., Zhi, J., Tian, T., Cihan, A., Cevher, M.A., Liu, Z., David, Y., Muir, T.W., Roeder, R.G., and Yu, M. (2021). DOT1L complex regulates transcriptional initiation in human erythroleukemic cells. Proc Natl Acad Sci U S A 118, e2106148118.

Yi, Q., Chen, Q., Liang, C., Yan, H., Zhang, Z., Xiang, X., Zhang, M., Qi, F., Zhou, L., and Wang, F. (2018). HP1 links centromeric heterochromatin to centromere cohesion in mammals. EMBO Reports 19, e45484.

Yu, W., Chory, E.J., Wernimont, A.K., Tempel, W., Scopton, A., Federation, A., Marineau, J.J., Qi, J., Barsyte-Lovejoy, D., Yi, J., et al. (2012). Catalytic site remodelling of the DOT1L methyltransferase by selective inhibitors. Nat Commun 3, 1288.

Zhang, Y., Liu, T., Meyer, C.A., Eeckhoute, J., Johnson, D.S., Bernstein, B.E., Nusbaum, C., Myers, R.M., Brown, M., Li, W., et al. (2008). Model-based analysis of ChIP-Seq (MACS). Genome Biol 9, R137.

Zikmund, T., Paszekova, H., Kokavec, J., Kerbs, P., Thakur, S., Turkova, T., Tauchmanova, P., Greif, P.A., and Stopka, T. (2020). Loss of ISWI ATPase SMARCA5 (SNF2H) in Acute Myeloid Leukemia Cells Inhibits Proliferation and Chromatid Cohesion. Int J Mol Sci 21, E2073.

Zofall, M., and Grewal, S.I.S. (2006). Swi6/HP1 Recruits a JmjC Domain Protein to Facilitate Transcription of Heterochromatic Repeats. Molecular Cell 22, 681–692.

